# Enhancer plasticity sustains oncogenic transformation and progression of B-Cell Acute Lymphoblastic leukemia

**DOI:** 10.1101/2022.05.18.492497

**Authors:** G Corleone, C Sorino, M Caforio, S Di Giovenale, F De Nicola, V Bertaina, A Pitisci, C Cortile, F Locatelli, V Folgiero, M Fanciulli

## Abstract

Growing evidence report that non-genetic-driven events such as enhancer reprogramming promote neoplastic transformation and strongly contribute to the phenotypical heterogeneity of cancers as much as genetic variation. In this context, we investigated the role of enhancers in sustaining oncogenic transformation in B-Cell Acute Lymphoblastic leukemia in children (BCP-ALL), a type of cancer caused by the accumulation of lymphoid progenitor cells in the bone marrow and a leading cause of cancer-related mortality in children. Using next-generation sequencing (ATAC-seq), we built the most up-to-date map of chromatin accessibility in pediatric BCP-ALL. We observed that enhancer activity dynamically changes during cancer progression and represents principal phenomena underlying phenotypic–functional characteristics of BCP-ALL progression. BCP-ALL patients are dominated by a regulatory repertoire (N=∼11k) originally represented at diagnosis that shrinks under treatments and subsequently re-expands, driving the relapse. We then deployed a wide range of in-vivo, in-vitro assays, and in-silico analyses to demonstrate the impact of enhancer activity in determining the phenotypical complexity. CRISPR-Cas-9-mediated validation of selected productive enhancers demonstrated a high capability of these regions to control MYB and DCTD oncogenic activities. Taken together, these findings provide direct support to the notion that enhancer plasticity is a crucial determinant of the BCP-ALL phenotype.

## INTRODUCTION

The major obstacle to effective cancer therapy is cancer intra- and inter-heterogeneity. Thus, the research of mechanisms that can act transversely across patient characteristics is of fundamental importance. A series of genetic events are commonly assumed to sustain the cellular substrate in the development of any cancer by supplying the oncogenic potential for proliferation and dissemination (1). However, genetic variegation does not entirely explain cell phenotypes (2,3). Indeed, recent analysis demonstrates that driver coding mutations rarely differentiate in metastatic samples despite the remarkable clinical and morphological differences then primaries (1,4). Nevertheless, it is well established that non-genetic-driven events, such as enhancer reprogramming, promote neoplastic transformation and strongly contribute to the phenotypical heterogeneity of cancers as much as genetic variation (1,5). In addition, pervasive transcription of genomic regions other than protein-coding genes generates tremendous non-coding RNA (ncRNA) species far more than previously recognized (6,7). Among these, RNAs generated from enhancers (i.e., eRNAs) have attracted particular interest due to their potential roles in mediating enhancer-gene interactions and their frequent overlap with disease-associated noncoding risk loci (8). Functionally, eRNAs can be considered an integral component of active enhancers, facilitating gene activation and/or enhancer-promoter loops by interacting with transcriptional activators and co-activators (9–13). Aberrant eRNA expression is highly associated with enhancer malfunction and is involved in dysregulation of oncogenes (14), tumor suppressor genes (15), as well as in abnormal cellular responses to external signals, such as hormone (16), inflammation (17,18) (19) and other stimuli (20,21). Numerous eRNAs are cancer type-specific and can be potentially employed for molecular diagnosis of cancer types with a significant prognostic utility. Indeed, data from 9000 sequenced tumors showed a marked prognostic significance of enhancer activation even higher than protein-coding genes. (22).

BCP-ALL is the most common malignancy in children, and while highly curable, the recurrence rate accounts for ∼10% of patients (23). Risk factors for outcome following the first relapse have been defined and incorporated into risk stratification schemes: time from diagnosis to relapse, site of relapse, immunophenotype, and minimal residual disease (MRD) response to reinduction therapy (23).

Little is known about the chromatin layout of BCP-ALL patients and the relative impact on tumorigenesis and drug response. One possibility is that chromatin plasticity affects the enhancer activity and engagement of TFs, thus supporting oncogenic pathways (24). Moreover, a comparison of chromatin openness variegation in primary, remission and relapse samples may provide insights into the biology of arising and response to therapy of cancer (24,25). Here we performed genome-wide profiling of the chromatin accessibility landscape of a longitudinal cohort of BCP-ALL cases at onset, remission and relapse. Using next-generation sequencing (ATAC-seq), we built the most up-to-date map of chromatin accessibility in pediatric BCP-ALL. Notably, these findings were summarized in a human primary cell line of BCP-ALL, associating enhancers activity with important target genes involved in disease transformation and progression. Finally, by CRISPR-Cas-9-mediated editing of selected eRNA sequences, we demonstrated the direct and effective control of the respective gene target expression. Altogether, these results reinforce the fundamental role of enhancers in the genesis and evolution of the BCP-ALL.

## RESULTS

### Genome-wide mapping of chromatin accessibility in a longitudinal cohort of pediatric BCP-ALL defines the disease stage

The cis-regulatory apparatus of BCP-ALL cells has never been described to date. We designed the study including 26 cases of BCP-ALL obtained at onset (N=11), remission (N=7), and relapse (N=8). To identify the diversity of malignant B cells we selected a control cohort of healthy bone marrow (HBM) from 6 age-matched or adult donors who donated bone marrow (BM) for transplantation. All patients were profiled for BCP-ALL most common molecular abnormalities using cytogenetics. Interestingly, 64% of diagnosed patients did not carry any genetic abnormality, while this frequency dropped to 25% at the relapse (Table 1). We calibrated our study by collecting B cells from fresh BM samples, isolating CD19+ cells by immunodensity from BM, and then profiling them by using ATAC-seq analysis. This strategy aimed to build a genome-wide map of accessible regions and infer the differential regulatory activity at accessible sites, ultimately gaining novel insights into BCP-ALL disease states. ATAC-seq experiments revealed accessible sites amongst all samples ranging between ∼20k to ∼80k sites, and our mapping strategy produced a cumulative number of 150,123 chromatin accessible sites (Fig.1a). Roughly 20% of these sites mapped within 5kb to the closest transcription starting sites (TSS) at loci putatively considered as promoters. Interestingly, all the selected groups of patients shared same proportion of promoter-like active sites (Extended Data Fig.1a). Notably, while healthy tissues were strongly defined by promoter like activity, the onset group linearly increased the number of active sites at distal loci (Extended Data Fig.1a). Furthermore, promoter activity was equally shared among our cohort (Extended Data Fig.1b-c) while the largest part of Cis-Regulatory Elements (CREs) was encoded at distal genomic loci accounting for ∼80% of the total number of sites. In addition, the onsets exhibit a much stronger activity at non-coding and intron regions than the other groups (Extended Data Fig.1b). Taken together our data confirm the notion that phenotypical heterogeneity of a cancer cell is defined by the setup of the distant regulatory asset (24).

**Fig.1:**
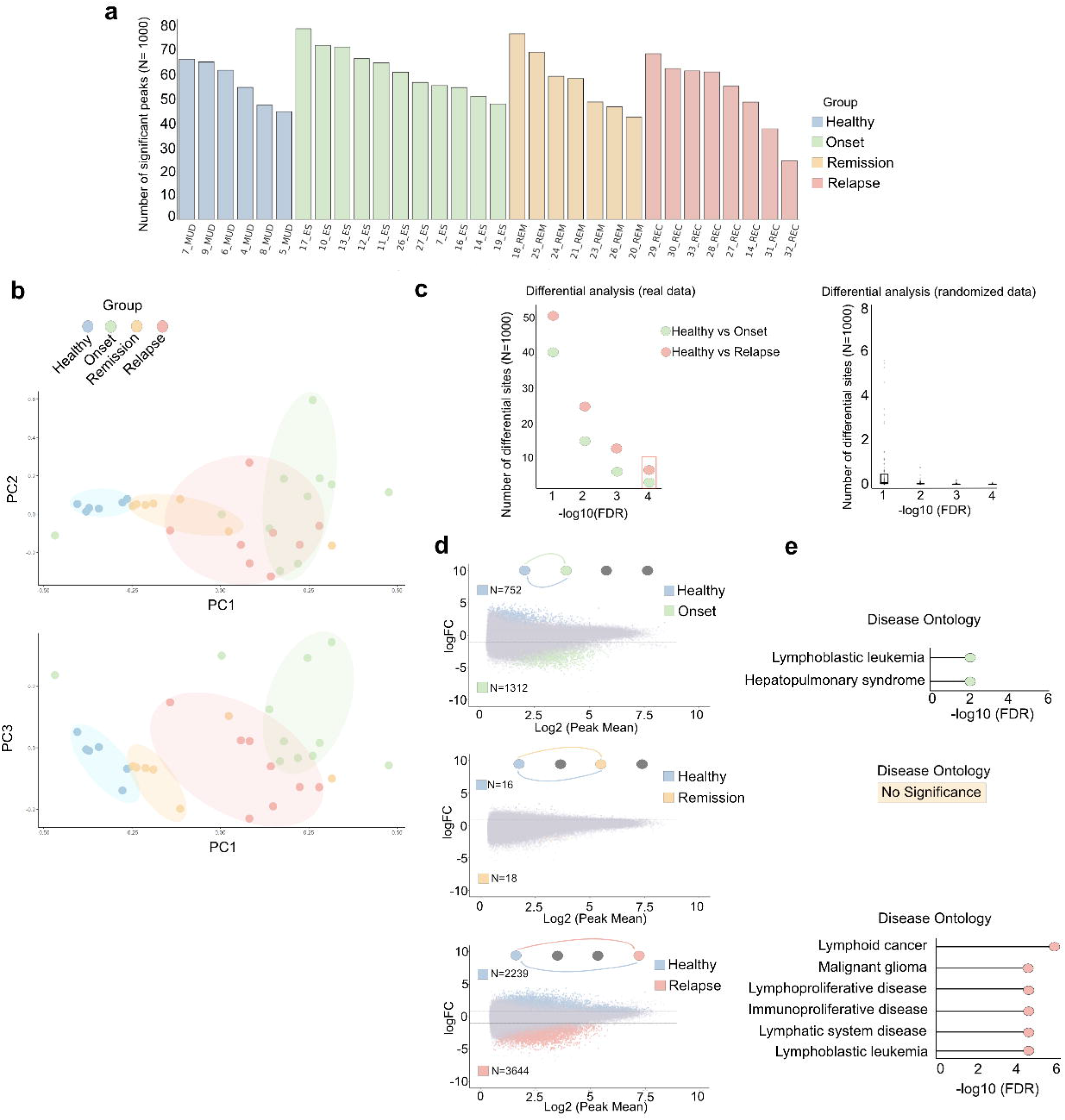
Differential analysis of BCP-ALL accessibility during cancer progression. **a**) Histogram showing the total number of ATAC-seq significant peaks x sample profiled. X-axis: name of the sample; Y-axis: Absolute number of significant peaks. Color legend: Blue= Healthy samples; Green= Samples at Onset; Orange= Samples at Remission; Relapse= Samples at relapse. **b**) PCA of accessibility profiles of our patient cohort. Up: PCA between principal component 1 (x-axis) and principal component 2 (y-axis). Down: PCA between principal component 1 (x-axis) and principal component 3 (y-axis). Shaded areas in the PCA plot represent 90% confidence ellipses. Color legend: Blue= Healthy samples; Green= Samples at Onset; Orange= Samples at Remission; Relapse= Samples at relapse. **c**) Left: Differential analyses of accessibility profiles between Healthy vs. Onset (green point) and Healthy vs. Relapse (red point). X-axis: different points of significance; Y-axis: Number of differential peaks identified in the analyses. Right: Differential analyses were performed by applying 100 random sampling from the patient cohort. Group sizes matched the Healthy, Onset, and Relapse cohort. X-axis: different points of significance; Y-axis: Number of differential peaks identified in the analyses. **d**) MA plot of the differential peak accessibility. Top: Healthy vs. Onset; Middle: Healthy vs. Remission; Bottom: Healthy vs. Relapse. X-axis: Log2(Peak mean), Y-axis: Log Fold Change of the differential accessibility of peaks. N= number of significant differential peaks identified in the analysis where logFC>0.7 is upregulation in Healthy; logFC<-0.7 is upregulation respectively at Onset (green), Remission (orange), and Relapse (red). **e**) Disease ontology associated with the differential analysis of accessibility. Top: Upregulation at the Onset; Middle: upregulation at the Remission; Bottom: upregulation at the Relapse. Analysis was performed against Healthy tissues. The analysis is performed with the GREAT tool.

Next, we tested the relationship between DNA accessibility landscape and BCP-ALL phenotype at different disease stages. Interestingly, we found that the chromatin accessibility landscape is a main determinant of patient clustering using Principal Component Analysis (PCA). Indeed, patients grouped accordingly to the disease stage. As expected, healthy and remission patients clustered homogenously and were very similar, while patients at onset and relapse were more heterogeneous, confirming how regulatory regions sustain distinct phenotypes in active cancer (Fig.1b).

### The cis-regulatory activity is a main phenotypical determinant of BCP-ALL

CREs are distant regulatory elements that are positively associated with gene transcription and thus key elements in determining cell phenotype (26,27). To study the process of CREs modulation during BCP-ALL progression, we surveyed the variability of each given 150,123 identified CRE by performing differential analysis amongst patients at healthy, onset, remission and relapse groups (Fig.1c). We found evidence of highly diverse epigenetic profiles in onset and relapse with at least 10-fold more differentially active CREs than expected by chance (Fig.1c, right). As previously observed in other cancer types (28), the prevalent differential patterns were exhibited between the healthy vs onset and healthy vs relapse groups, while remissions showed only a marginal number of differential sites than healthy (Fig.1d and Extended Data Fig.1d). By applying a very stringent threshold of FDR<10^-4^ of significance, we selected 1312 CRE sites upregulated in the onset which strikingly, enriched with ontologies specifically associated with Lymphoblastic Leukemia (Fig.1e). Conversely, differential analysis between healthy and remission showed a recovery of the healthy-like accessibility asset after therapy with only 34 CREs significantly modulated at the remission. Furthermore, we obtained the largest diversification of CREs at the healthy-remission vs relapse analysis (Fig.1d, Extended Data Fig.1d-e). A totality of 5883 sites were actively modulated of which 3644 upregulated in the relapse (Fig.1d bottom). The interrogation of disease ontology showed a more robust enrichment of Lymphoid cancer disease and Lymphoblastic disease, suggesting a post therapy selection of CREs associated to the cancer phenotype (Fig.1e, Extended Data Fig.1f). Overall, these data support the hypothesis that BCP-ALL is sustained by plastic activity of CREs which strongly determine the post treatment behavior of cells.

### Dissection and tracking of cis-regulatory element activity during BCP-ALL evolution

The dynamics of cis-regulatory activity during the BCP-ALL span have never been comprehensively investigated to date. The use of epigenetic modifications to annotate CRE activity has been recently successfully applied (29). Indeed, increasing evidence suggests that chromatin accessibility identified by ATAC-seq is strongly linked to CRE activity (30,31). In addition, single-cell studies showed that bulk ATAC-seq and histone mark signals (H3K27ac and H3K4me3) are proportional to the cell contributing to it (32,33). Thus, each nucleosome positioning can be inferred as digital information where the state on/off directly corresponds to the single-cell state (34). This suggests that each bulk ATAC-seq signal directly relates to its relative clonality within the sequenced sample. Applying this concept to a phenotypically homogeneous population of cells for each sequenced samples would provide opportunities to deepen the understanding of the clonality of each CRE at any given sample. On this basis, we reasoned that a bulk ATAC-seq signal would be an effective tool for identifying CRE activity from a cohort of 100% pure CD19+ lymphocytes obtained from BCP-ALL patients. Following the strategy recently applied in solid tumors (34) we assigned a clonality score to each putative regulatory region identified by ATAC-seq in our patient cohort, coupled by a penetrance score which reflects the number of patients in the cohort sharing the activity of any given region (Fig.2a). This strategy allowed the dissection of the cis-regulatory activity of any given CRE in our longitudinal patient cohort. Each CRE region received assigned a score of clonality and penetrance. We then developed a strategy to identify CRE which were significantly modulated during the disease focusing on regions matching these criteria: 1) being positively modulated during the passage from healthy to onset, 2) being negatively modulated during onset and remission, and 3) being positively regulated between remission to relapse. Then, the selected regions were further integrated into a multistep process including: 1) integration with the most up to date pan-cancer data to infer high fidelity enhancers and super-enhancers, 2) integration with data obtained from an onset BCP-ALL cell line profiled with H3K27ac ChIP-seq, ATAC-seq, RNA-seq, Promoter-capture-seq followed by experimental validation including CRISPR-cas KO of the selected CREs (Fig.2a).

**Fig. 2:**
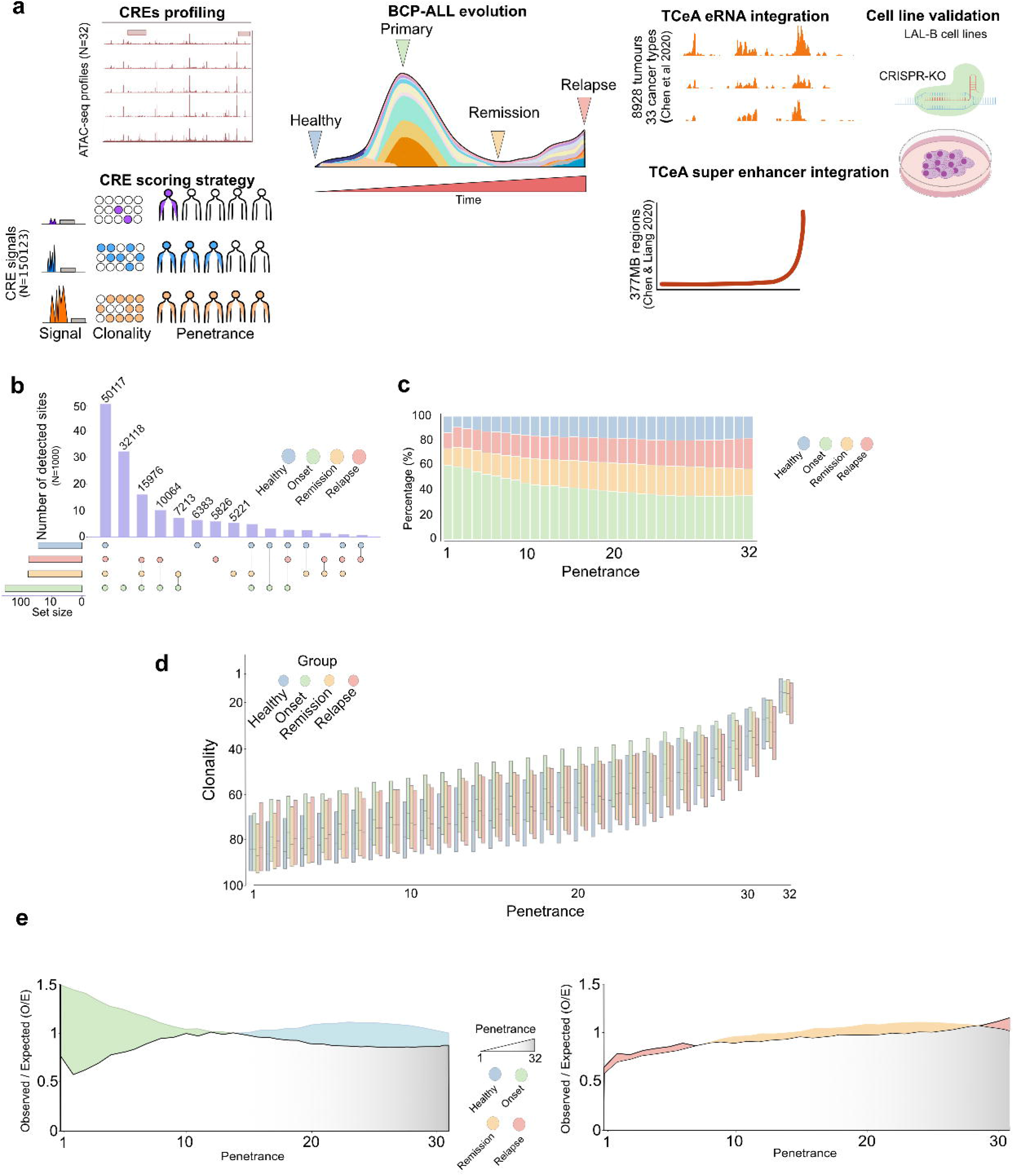
Dissection of cis-regulatory heterogeneity of BCP-ALL. **a**) Workflow of the study. From left to right: We profiled 32 samples of BCP-ALL to identify putative cis-regulatory regions. With a scoring strategy based on the Clonality and Penetrance indices, we dissected the accessibility landscape and prioritized the study toward the most clonal/penetrant cis-regulatory sites. Then, these two scores assigned to each CRE were used to monitor clonality and penetrance during BCP-ALL evolution by assessing the variation of CREs modulation at Healthy> Primary> Remission > Relapse stages. To provide more insights into the functional role of the selected CREs, we integrated data from the TCEA portal (cit), which provide enhancer RNA-seq profiles from 8928 samples of 33 cancer. Furthermore, we integrated a 377MB region of super-enhancer into our selection. We validated several elements with CRISPR KO and experimental procedures among all the selected CRE sustaining BCP-ALL progression. **b**) Upset plot of detected peaks among the different groups of patients. X-axis: Intersection combination; Y-axis: the absolute number of detected sites. Color legend: Blue= Healthy samples; Green= Samples at Onset; Orange= Samples at Remission; Relapse= Samples at relapse; Violet: Barchart of the number of detected sites at each intersection. **c**) Left: Stacked bar chart representing the percentage of significant peaks (x-axis) x group in the function of the penetrance index (x-axis); Right: Stacked bar chart representing the percentage of peaks x groups in the function of the distance to the closest annotated Transcription Starting Site in hg19 reference genome (TSS). Color legend: Blue= Healthy samples; Green= Samples at Onset; Orange= Samples at Remission; Relapse= Samples at relapse. **d**) Boxplots show the median Clonality Index value and interquartile ranges for each detected peak x disease stage in the function of the Penetrance index. Color legend: Blue= Healthy samples; Green= Samples at Onset; Orange= Samples at Remission; Relapse= Samples at relapse. **e**) Observed/Expected (O/E) ratio of peaks (y-axis) at any penetrance score (x-axis) between Healthy vs Onset (left) and Remission vs Relapse (right). Color legend: Blue= Healthy samples; Green= Samples at Onset; Orange= Samples at Remission; Relapse= Samples at relapse.

### The accessibility landscape of BCP-ALL

We tested whether the identification of CREs was associated with cancer-specific stages. First, we classified the repertoire of the regulatory potential at each stage separately. Secondly, we annotated the CREs detected and then, calculated the absolute difference of activated regulatory repertoire in the groups. Not surprisingly, the onset group enclosed the highest number of activated CREs, accounting for more than 120,000 detected sites (Fig. 2b). Roughly 50k CREs were shared among each disease stage, while 32,118 CREs were present only in the onset group. A subset of 15,976 CREs was detected in leukemic samples but not in the healthy samples (Fig.2b). Among all sites, 60% were only detected as private or low penetrant in the onset group. This percentage fell to 35% in more penetrant regions of the cohort. On the other hand, the healthy, remission, and relapse group exhibited less than 10% of detected sites at private or low shared regions, suggesting that the onset group is characterized by a larger plethora of distinct cell subclones than the other groups (Fig.2c).

Next, to provide qualitative insights into the relative contribution of each detected CREs to the BCP-ALL phenotype, we applied a computational framework that allows dissecting the regulatory heterogeneity of the chromatin accessibility landscape. We observed that penetrance and clonality were in strong linear relationship in each sample group (Fig. 2d). Of note, the onset group exhibited systematically higher clonality of CREs than the other groups in the function of the penetrance index, thus suggesting a larger engagement of sample-specific regulatory potential. These observations were further corroborated by a linear regression analysis independently performed in each disease group, demonstrating a positive relationship between clonality and penetrance (Extended Data Fig.2a). Next, the stratification of the chromatin accessibility landscape conferred an advantage in evaluating the significance of the heterogeneity seen in the onset in relationship with the other groups. Notably, the onset group exhibited higher observed heterogeneity than expected at low penetrance indexes (PI=1-14), while highly penetrant CRE were observed more in healthy samples than expected (Fig.2e). On the other hand, the remission to relapse stage did not significantly change.

Taken together, these findings provide direct support to the notion that plasticity of the regulatory elements is a key determinant of cancer-specific phenotype, and sub-clonal diversification of cancer-specific sites occurs at the passage between a healthy state to the onset to then recover to a healthy-like layout after treatment.

### The regulatory landscape of BCP-ALL dynamically changes during cancer evolution

We carried out a phenotype-driven computational analysis to investigate the regulatory variability of chromatin openness in patients during BCP-ALL evolution. We hypothesized that a subset of regulatory regions was dynamically selected to drive the Onset, Remission, and Relapse group phenotype. To test this, we selected highly penetrant regulatory sites driving cancer (Fig.3a), showing poor activity in the healthy tissues, dynamically changing their relative clonality over time. After tracing the behavior of 150,123 CREs, our strategy identified 11,083 CREs exhibiting a significant change in clonality, with a marked increase from healthy to onset, then a drop after therapy and again an increase from remission to relapse (Fig.3a). In agreement, unsupervised clustering of normalized ATAC-seq enrichment at the selected CREs highlighted a surprising similarity between the healthy and remission groups profiles except for two samples at the remission, which were clustering to the onset and relapse groups. Hierarchical clustering of the relative enrichment of each CRE determined two major clades, one including the higher activity of CREs at the onset and relapse (C1 and C2), and one with an enrichment higher in healthy and remission (C3 and C4) (Fig.3b). The selected CRE were largely penetrant at the onset and private in healthy and remission tissues while more heterogeneous at relapse (Extended Data Fig.3a). Of note, ∼90% to the selected CREs were distal to the closest gene (Extended Data Fig.3b). Functional characterization demonstrated that our approach successfully targeted regulatory sites strongly involved in lymphocyte activation/differentiation and in sustaining the regulation of genes linked to lymphoblastic leukemia, lymphadenopathy, and more generally, auto-immune diseases (Extended Data Fig.3c). Moreover, since the accessibility of CREs is largely controlled by transcription factors (TFs), we investigated the most enriched binding motifs in our selected sites to identify their cognate TFs (Fig.3c). We inferred the TFs putatively binding to the selected CREs by performing TF motif analysis. Then, we calculated the observed/expected ratio of the most significant in the first clade (C1 and C2 clusters) of sites. Interestingly, the top significant identified TFs were well-known drivers of B cell development with established oncogenic potential. Indeed, numerous studies have already reported that aberrant modulation of EBF1, ETS1, ERG, RUNX transcription factors have profound effects on lymphoid neoplasms derived from B cell progenitors (35,36). In sum, this analysis indicates that BCP-ALL progression is associated with a plastic reprogramming of the non-coding regulatory landscape of cells which involves the recruiting of key regulatory elements of hematopoiesis.

**Fig. 3:**
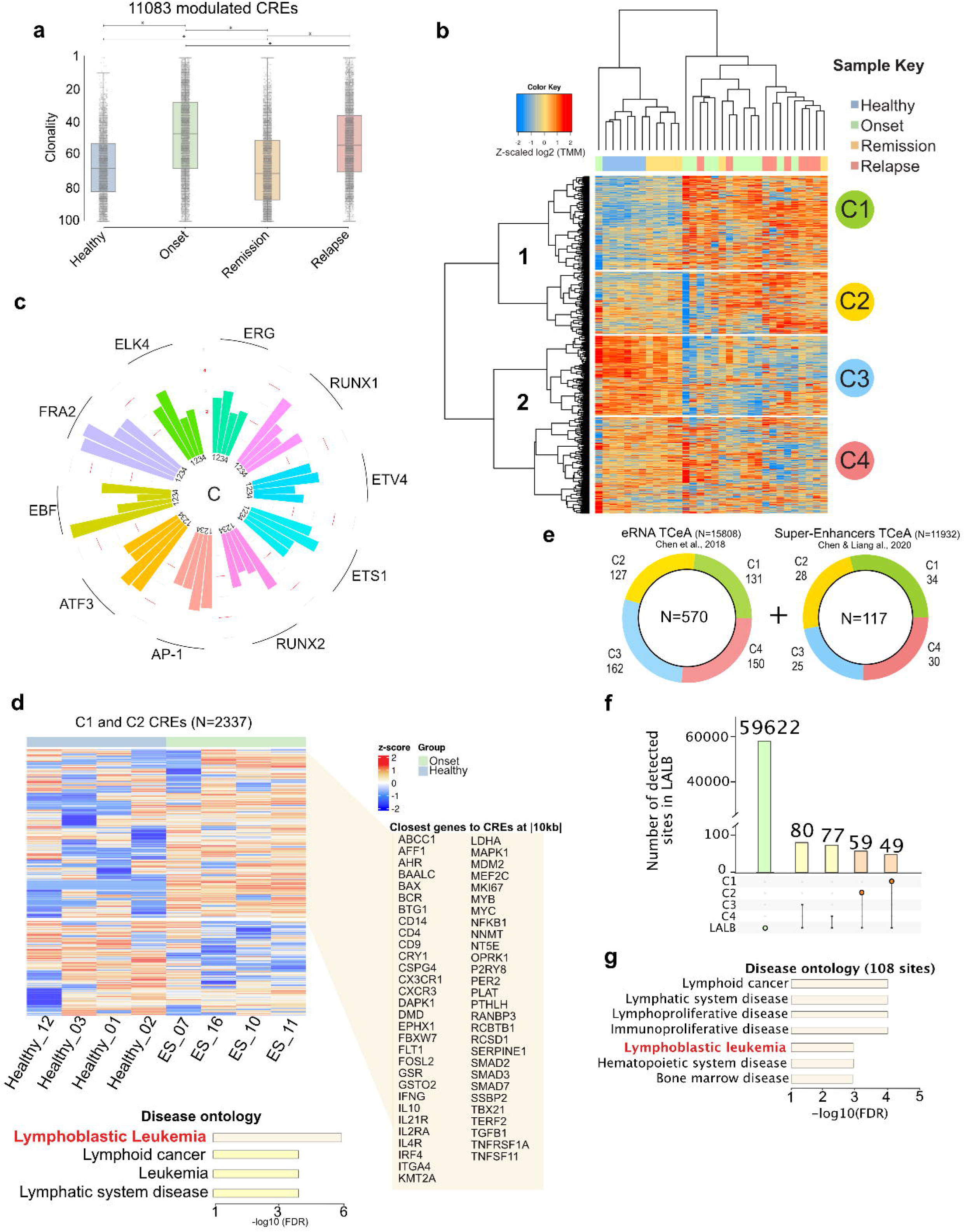
Enhancer engagement varies amongst BCP-ALL cancer stages. **a**) Boxplot depicting the Clonality index of the selected CREs at the Healthy, Onset, Remission, and Relapse status. Color legend: Blue= Healthy samples; Green= Samples at Onset; Orange= Samples at Remission; Relapse= Samples at relapse. Statistical test: Kruskal-Wallis test followed by Dunn’s test. Pval= *<10^-^4. Statistical significance was calculated using a pairwise, two-tailed t-test. **b**) Unsupervised Clustering Heatmap showing z-scaled log2(TMM) score enrichment of the selected CREs (N=) in each given patient of the cohort. The analysis identified two main branches of data (left) and four main clusters named C1, C2, C3, and C4 (right). Color legend: Blue= Healthy samples; Green= Samples at Onset; Orange= Samples at Remission; Relapse= Samples at relapse. **c**) Polar bar plot depicting transcription factor motif enrichment at C1, C2, C3, and C4 sites. Bar plot representing the Observed/Expected ratio of the transcription factor motif at any given C cluster. **d**) Unsupervised heatmap depicting eRNA at selected CREs (C1, C2) not matching exons annotation (hg19). RNA-seq data from the sample cohort composed 4 Healthy tissues and four tissues at the Onset. Data were normalized and scaled with z-scoring. The window (right) highlights the closest genes (distance ranging |10kb| from each CREs) associated with the upregulated regions at the Onset. Disease Ontology (bottom) of upregulated CREs at the Onset obtained with GREAT tool. Color legend: Blue= Healthy samples; Green= Samples at Onset E) Stacked pie chart of C1, C2, 3, C4 selected CREs intersected with ERNA TCeA portal (left) and Super-Enhancers TCeA portal (Right). Color legend: Green= C1; Yellow= C2; Blue= C3; Red= C4. **f**) Upset plot of C1, C2, C3, and C4 selected CREs and LALB ATAC-seq peaks. X-axis: Intersection combination; Y-axis: the absolute number of detected sites at each intersection. **g**) Disease ontology of 108 CREs in C1 and C2 matching LALB cells. Ontology obtained with GREAT tool.

### Active enhancers are key elements of BCP-ALL progression

The functional characterization of a given CRE is an inherently challenging process. This is due to the complexity of the molecular events, the variability in the addiction of TFs, and the elusive cross-talks among known and unknown factors that orchestrate gene expression in the given time (37). The lack of general rules in identifying and constructing the link between CREs and gene target signals encouraged us to apply a conservative approach by focusing on productive enhancer elements. Thus, we performed RNA-seq of 4 healthy, 4 onsets and measured the productivity at CREs loci which showed upregulation in the previously identified as C1 and C2. Unsupervised hierarchical clustering showed marked productivity of eRNA in 1092 CREs in the onset than healthy samples (Fig.3d top). We then linked these productive elements to the closest gene and enrichment analysis strongly enriched to lymphoblastic leukemia (FDR< 10^-6^) and Lymphoid cancer (FDR< 10^-3^) (Fig.3d bottom). The selected CREs were potentially regulating genes previously identified as key determinant of BCP-ALL phenotype such as ERG, KMT2A or MYB (38,39). To build a more accurate classification of the selected productive enhancer-like CREs, we integrated our analysis with pre-annotated productive enhancer and superenhancer (40) together with the accessibility landscape of LAL-B, a primary cell line generated by manipulation of BM from patient at BCP-ALL onset (41). In line with this, 570 enhancers and 117 super enhancers were selected among the C1, C2, C3, C4 clusters (Fig.3e). In addition, we noticed that the number of putatively eRNA productive sites at each given sample of our cohort was highly heterogeneous and that peaks at the onset were generally more clonal (Extended Data Fig.3d). Then, we compared LAL-B accessibility profiles with the patient cohort (Extended Data Fig. 3e) and observed 108 active CREs shared with C1 and C2 (Fig. 3f), which were linked to key determinants of lymphoid and bone marrow neoplasms among which lymphoblastic leukemia was found (FDR<10^-3^) (Fig.3g). Then, we measured the binding of the identified TFs observing that RUNX2, ERG and ETS1 may bound to over then 75% of the selected loci (Extended Data Fig.3f). We ultimately measured the pan-cancer cell line profiles of H3K27ac at the selected loci (Extended Data Fig.4a) showing a large variegation of enhancer activity not specifically lymphoid dependent. Overall, these data demonstrate that a subset of productive and plastic enhancers sustains BCP-ALL phenotype.

### Long-range chromatin interactions add functional insights into BCP-ALL primary cell line

Distal regulatory elements function by physically interacting with target genes through chromatin looping. However, predicting enhancer-gene interaction in a given cell type context lacks general rules that can be uniformly applied (42). Therefore, our primary effort was to determine the univocal genes regulated by the selected enhancers. To address the functional role of the selected CREs, we first profiled the chromatin interacting landscape of LAL-B cells, using in situ-promoter capture HiC (43). This analysis revealed short and long-range interactions by analyzing the data at three different map resolutions (5kb, 10kb, 25kb) (Fig.4a). Our analysis detected 30190 genomic interactions, of which ∼15k were classified as a promoter – CRE looping. About 11k loops were detected between 2 non-promoter CREs, while 3745 loops were observed amongst two known promoters (Fig.4a, left). Interestingly, the detected Promoter-CREs interactions were observed at a distance between 50 to 500kb. Only a minority of loops were detected at a distance of more than 1000kb from the promoter. Nevertheless, we identified only ∼1000 interactions at a distance shorter than 50kb (Fig.4a, right). This suggests that promoter capture HiC not only identifies loops among a known promoter with unknown distal CRE but also captures the full spectrum of superimposed folding states and interactions among non-promoter CREs. For example, at the MYC locus, we observed more than 20 direct interactions Promoter-CRE and 12 undirect interactions generated by multiple looping towards different CREs that ultimately connect to the MYC promoter (Extended Data Fig.5a). Although recent data confirm that the enhancer gene-target prediction can be inferred simply by gene proximity at high precision (44), we integrated our Promoter-capture HiC approach with ATAC-seq results from patients and LAL-B cell line, and CTCF and Pol2 ChIA-Pet of K562 cell line to survey every possible enhancer-gene contact (Fig.4b, Extended Data Fig.5a) (Table 2). Then, we measured the transcriptional and accessibility output (Fig.4b, Extended Data Fig.5b) of the selected gene targets in LAL-B cells together with lymphoid cells at three differentiation steps (Naïve B-cell, Mem-B cell, and Plasmablast) obtained by healthy individuals (45). Applying this strategy, we observed 106 genes exhibiting a marked transcriptional output specific to only the LAL-B cells. The selected genes list included genes previously implicated to the B type ALL and hematopoietic malignancies such as EBF1, MYB, ETS1 and MYC (35,36). (Fig.4b). Next, we sought to quantify and characterize the essentiality of the selected genes in the recently screened genome-scale CRISPR–Cas9 loss-of-function pediatric cell lines available in the Dependency Map Portal (DEPMAP) (46). Thus, we measured the essentiality score (47) of each gene in the whole set of screened cell lines (N=∼1000 cell lines) which include 11 B-ALL cell lines (Extended Data Fig.5c). We selected the top 100 most dependent cell lines of each given gene and counted the number of B-ALL cell lines included in the selection (Fig.4c) to identify the genes affecting more specifically BCP-ALL fitness than others, observing that two genes were exhibiting a marked specify to ALL-B cell lines viability together with a physical enhancer-promoter interaction: DCTD, MYB (Fig. 4c-d).

**Fig. 4:**
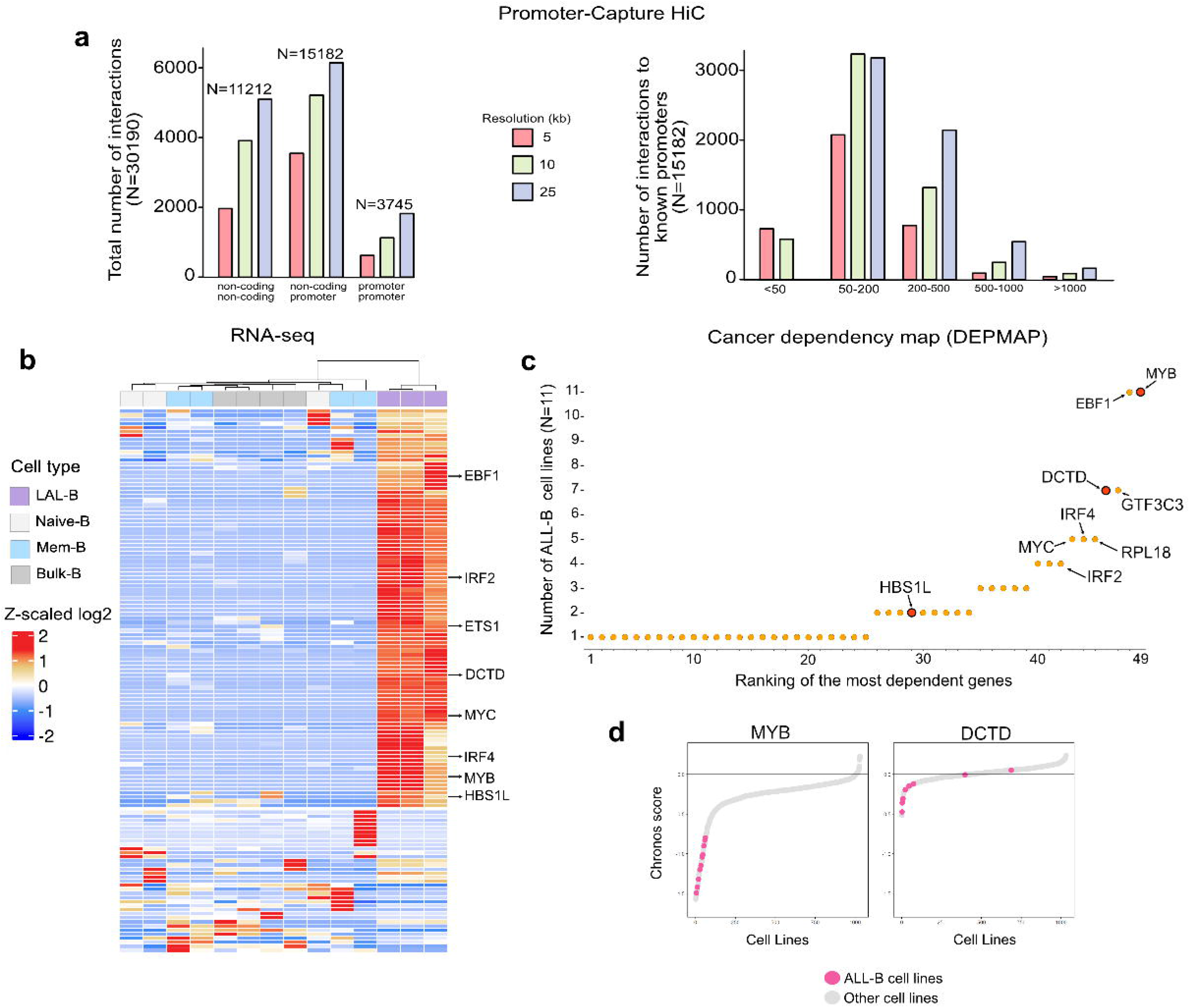
Identification of enhancer-target genes in LAL-B. **a)** Barchart summarizes the number of absolute significant interactions of Promoter-Capture HiC in LALB cells grouped by annotation of anchor and target in the genomic context (left) and the number of absolute interactions specific to only CRE-Promoters (hg19) in LALB grouped by range distance (right). Color legend: Red= 5kb resolution; Green= 10kb resolution; Violet= 25kb resolution. **b**) Unsupervised clustering of gene expression (RNA-seq) of LAL-B, Naïve-B, Mem-B, and Bulk-B cells (triplicates for each category) obtained from Calderon et al. Nat. Gen 2019. Data were normalized and scaled with z-scoring. Genes interrogated in the heatmap (N=) are selected by evidence of looping with the 108 selected CREs. Color legend: Violet= LAL-B cells; Light grey= Naïve-B; Blu= Mem-B; Grey=Bulk-B cells. **c**) Plot showing the ranked most dependent genes of BCP-ALL among the selected from figure 4B (x-axis) in the function of the number of ALL-B cell lines (N=11) ranking at the top 10% of the most sensitive cell lines among the total number of available cell lines in the DEPMAP portal at any given gene. **d**) Plots showing the Chronos score of MYB and DCTD genes of the selected ALL-B cell lines (pink) and all the other cell lines (grey).

Collectively, these data show that our strategy captures qualitative properties in BCP-ALL evolution. DCTD and MYB enhancer activity marks a dominant phenotypic clone specific to the disease process and to its treatment response.

### MYB de novo activated enhancers are important regulatory elements in BCP-ALL

The transcription factor MYB plays a key role in regulating hematopoiesis (48). Single-cell data show that MYB is highly expressed in the myeloid development and epithelial cells while transcriptionally silent in most lymphoid cells (Extended Data Fig.6a). However, MYB dysregulation is often associated with myeloid and lymphoid hematological neoplasms, including acute lymphoblastic leukemia (49,50). In addition, several solid tumors show an aberrant expression of MYB (50) (Extended Data Fig. 5b). Of note, qRT-PCR analysis revealed increased expression of this gene in Onset and Relapse compared to Healthy and remission samples (Extended Data Fig. 5c).

The main alterations underlying the increased expression of MYB in tumors are caused to chromosomal translocations, gene duplications, or juxtaposing active enhancers from other genomic regions (51). Recent studies report that MYB expression is tuned by activity of a CREs cluster dwelling in the intergenic region spanning the 135 kb between MYB and HBS1L genes (52–54). This region contains numerous enhancer elements capable of looping towards the MYB promoter and regulating its relative gene expression. (52–54). In particular, genome-wide association studies (GWAS) report that SNPs within this region affects the normal development of erythrocytes and platelets (55–57), and these SNPs are associated to sickle cell disease and B thalassemia. Subsequent studies have shown that enhancers at -84 and -71 to the MYB promoter are critical regulators of erythropoiesis (57). More recently, enhancers activity at -38 and -88 to MYB promoter has been associated with human myeloid leukemia cell lines (54). Interestingly, ATAC-seq single-cell analysis of plasma cell and memory B cells shows that the enhancer cluster is not accessible while both HBS1L and MYB promoters are in the active state (Extended Data Fig.5d). These observations suggest that although MYB gene expression is fundamental in developing healthy myeloid cells, it is aberrantly regulated in different cancers.

Our strategy has identified two enhancer elements within the 135kb MYB-HBS1L region, affecting BCP-ALL phenotype. Accordingly, we performed a comprehensive analysis to characterize the relationship between the selected enhancer and MYB regulation in the BCP-ALL context. We first collected healthy and BCP-ALL samples at onset, remission, and relapse (N=17 samples) and evaluated MYB protein expression (Fig.5a). Our data clearly show that MYB protein significantly emerges in the onset and the relapse samples while dramatically reducing in a healthy-like state after treatment. Promoter capture Hi-C sequencing confirmed that the selected elements loop toward MYB and HBS1L genes (Fig. 5b). ChIA-Pet data in the K562 cells corroborated the hypothesis that the 135 kb MYB-HBS1L intergenic region comprises numerous CREs interacting with both genes. These enhancers in our cohort resulted nucleosome depleted and eRNA productive (Fig. 5c). Qualitative analysis showed that 51kb and 67kb enhancers are rarely clonal in healthy tissues. The clonality score significantly increases in the passage from healthy to onset state. Then, they resulted differentially harmed by the therapy. Enhancer at 51kb was silenced in 50% of the remissions but was clonal in the relapse. The enhancer at 67kb was refractory to the therapy and showed high clonality in 100% of the considered samples (Fig. 5c). In addition, we measured the ATAC-seq ratio between each enhancer and MYB promoter, showing the marked increase of enhancer clonality in the disease stages than in healthy samples (Extended data Fig. 6e). Notably, despite being crucial in myeloid leukemia, enhancer at 38kb was active in only a tiny proportion of patients and active only in one of our relapse samples. These data prompted us to further characterize these two regions by using the CRISPR-Cas9 strategy. We found that inactivation of both these two distal enhancers significantly reduced their eRNA transcripts (Figs.5d left and 5e left). While the inhibition -51 kb region negatively reduced both the protein and gene production of MYB and HBS1L, the inactivation of the -67 kb region was able to inhibit only MYB gene (Figs. 5d middle, 5e middle and Extended Data Figs. 6f). Importantly, these results were associated with a significant reduction of growth rate as compared to control cells (Figs. 5d right, 5e right). Collectively, our results identify two new selectively activated enhancers in BCP-ALL, which significantly modulate MYB expression and tumor cell proliferation together with BCP-ALL growth and progression.

**Fig. 5:**
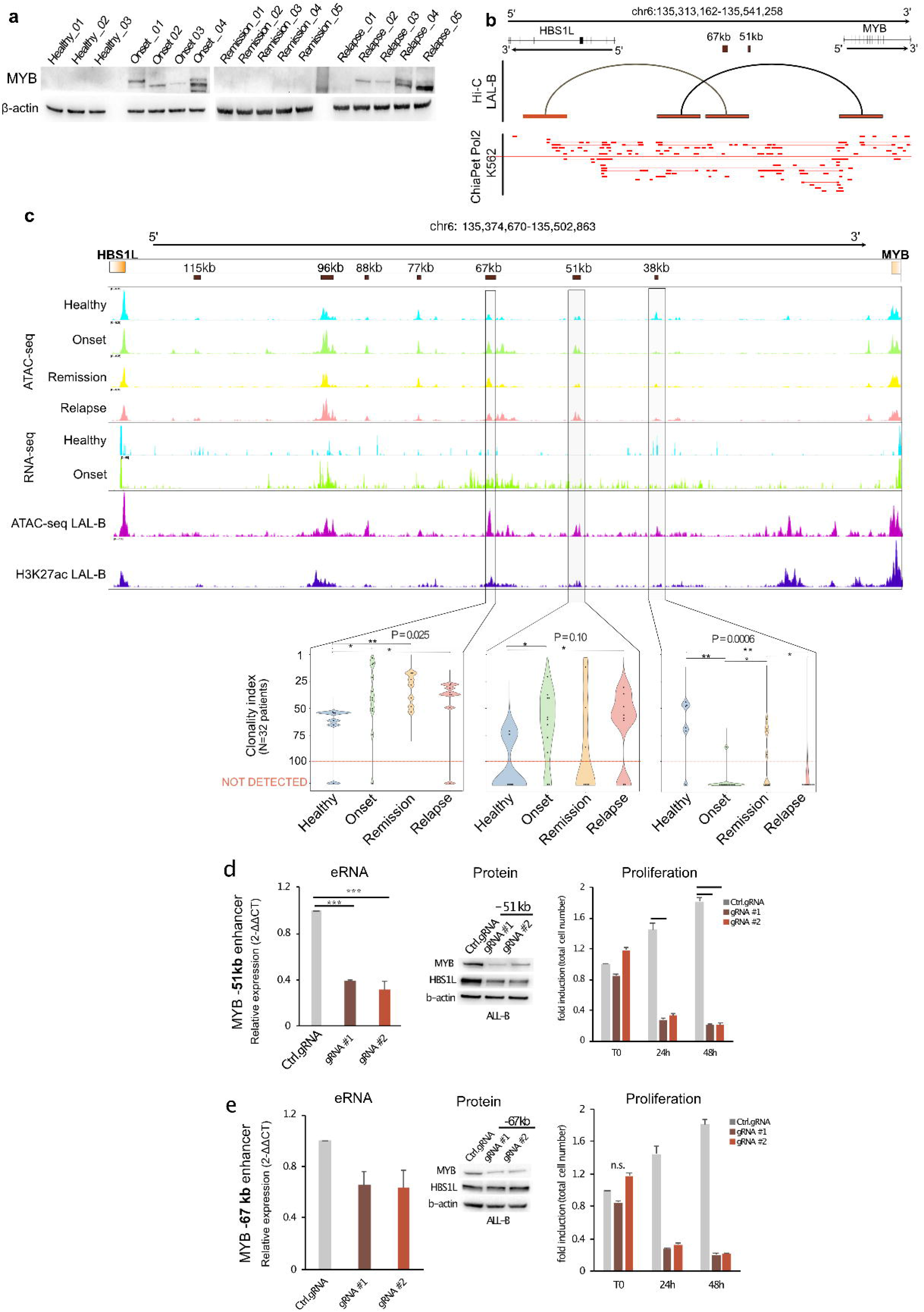
Myb enhancers sustain BCP-ALL progression. **a**) Relative expression of Myb protein determined by Western Blot (WB) in healthy (N=3), onset (N=4), remission (N=5), and relapse (N=5). β-actin was used as loading control. **b**) Chromatin looping identified by Hi-C Promoter-Capture sequencing at the MYB/HBS1L genomic window (top), integrated with Pol2 ChIA-PET data of K562 downloaded from ENCODE. Selected CRE elements are depicted in the dark brown boxes. Red boxes show looping genomic interactors identified by JUICER. **c**) ATAC-seq, RNA-seq profiles of our patient cohort at Healthy Onset, Remission, Relapse (ATAC-seq) and at Healthy and Onset (RNA-seq) at the MYB/HBS1L genomic window, ATAC-seq of LAL-B and H3K27ac ChIP-seq of LAL-B. Black boxes show the identified CREs within the window. Light grey windows highlight selected CREs experimentally validated. Together with violin plots depicting the Clonality index of the given CRE in the patient cohort. Pval represented at the top of each violin plot group is obtained by applying the Kruskal-Wallis chi-squared The statistical test applied: Pairwise Wilcoxon rank-sum test. *= Pval< 0.05. Color legend of violin plot: Blue= Healthy samples; Green= Samples at Onset; Orange= Samples at Remission; Relapse= Samples at relapse. **d**) Left, quantitative RT-PCR (qRT-PCR) analysis for Myb expression performed in B-ALL cells following CRISPR/Cas-9 of -51 kb region using two different gRNAs (#1-#2), compared to a control gRNA. Relative fold changes were determined by the comparative threshold (ΛΛCt) method using β-actin as endogenous normalization control. Data are presented as mean ± SD of three independent experiments: middle, WB with the indicated antibodies in control gRNA and -51 kb gRNA #1 and #2. β-actin was used as loading control; right, cell number analysis of control cells and -51 kb depleted cells at different time points. Data are presented as mean ± SD of three independent experiments. **e)** Left, qRT-PCR analysis for Myb expression performed in B-ALL cells following CRISPR/Cas-9 of -67 kb region using two different gRNAs (#1-#2), compared to a control gRNA. Relative fold changes were determined by the comparative threshold (ΛΛCt) method using β-actin as endogenous normalization control. Data are presented as mean ± SD of three independent experiments; middle, WB with the indicated antibodies in control gRNA and -67 kb gRNA #1 and #2. β-actin was used as loading control; right, cell number analysis of control cells and -67 kb depleted cells at different time points. Data are presented as mean ± SD of three independent experiments. **P≤ 0,01, ***P≤ 0,001 by Student’s *t*-test.

### DCTD expression is regulated by a distal element in BCP-ALL

Our study identified a critical enhancer regulating the Deoxycytidine monophosphate deaminase (DCTD) gene expression. DCTD is a key enzyme in the synthesis of genetic material which catalyzes the deamination of dCMP to dUMP, the nucleotide substrate for thymidylate synthase (58). Since its fundament role, the DCTD gene is ubiquitously expressed in all human healthy and neoplastic cells (Extended Figs. 7a, 7b and 7c); however, its relative role in cancer is controversial and partially understood. Indeed, TCGA and GETx bulk RNA-seq data prove that DCTD expression largely varies among different cancers (Extended Fig. 6c). Interestingly, DCTD is overexpressed in malignant gliomas (59), and DLBC (60) and marked as an adverse prognostic factor. In addition, DCTD has long been associated with chemoresistance to gemcitabine (61,62). Conversely, the DCTD gene is markedly downregulated in Acute Myeloid Leukemia (LAML) (Extended data Fig. 6c). Notably, the Cancer Dependency Map (Dep-Map) database revealed that DCTD KO rarely has an impact on cell proliferation, and only B-lymphoblastic leukemia cells are affected by the DCTD depletion (63) (Figs. 4c, 4d and Extended data Fig. 5c). Data from our cohort showed that the expression of the DCTD gene was strongly increased in BCP-ALL patients both at the protein and RNA level (Fig.6a and Extended Data 7e), specifically at the onset and the relapse stage. DCTD depletion by siRNA affected primary LAL-B cell proliferation (Extended Data 7f). ATAC-seq profiling of our patient cohort identified a 108 kb distant region to the DCTD promoter (Fig.6c). Promoter capture Hi-C in LAL-B cells confirmed that this region physically interacts with the DCTD promoter (Fig. 6b). Strikingly, this region was found to be much more accessible in leukemic patients than in healthy controls, suggesting its involvement in increasing DCTD expression in this disease (Fig.6c.). In addition, we demonstrated that the identified region was dynamically modulated in dependency on the disease stage (Fig.6c, Extended Data Fig.7g). Then, we applied CRISPR KO to the selected regulatory region showing significant downregulation of eRNA and DCTD RNA production, followed by protein clearance and lower cell proliferation capacity (Fig. 6d).

**Fig. 6:**
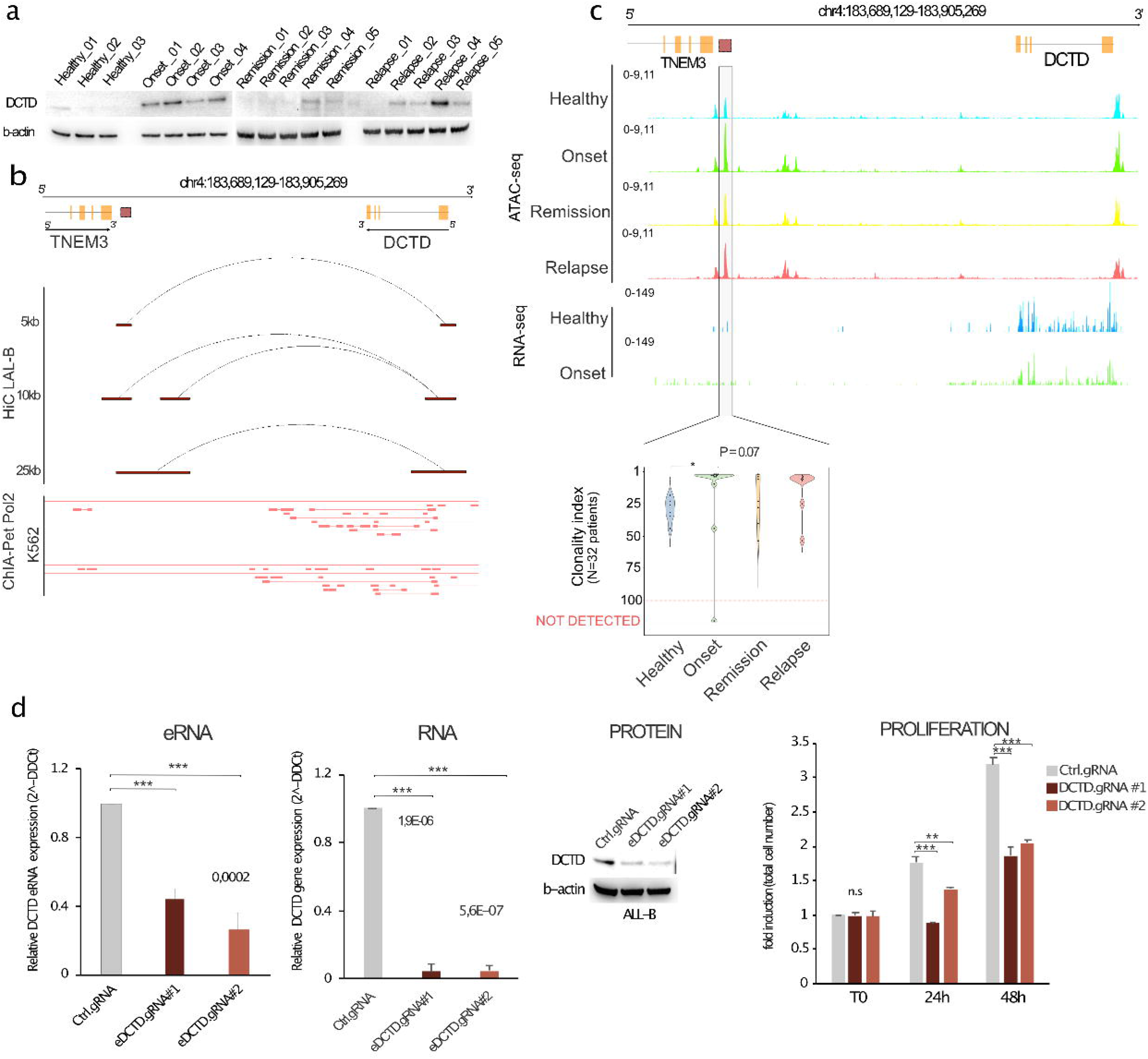
DCTD enhancer is a dominant clone of BCP-ALL progression. **a**) Relative expression of DCTD protein determined by WB in Healthy (N=3), Onset (N=4), Remission (N=5), and Relapse (N=5). β-actin was used as loading control. **b**) Chromatin looping identified by Hi-C Promoter-Capture sequencing at the TNEM/DCTD intergenic region (top), integrated with Pol2 ChIA-PET data of K562 downloaded from ENCODE. Selected CRE element is depicted in the dark brown box. Red boxes show looping genomic interactors identified by JUICER at different resolution. **c**) ATAC-seq, RNA-seq profiles of our patient cohort at Healthy Onset, Remission, Relapse (ATAC-seq) and at Healthy and Onset (RNA-seq) at the TNEM/DCTD genomic window. Black boxes show the identified CREs within the window. Light grey windows highlight selected CREs experimentally validated. Together with violin plots depicting the Clonality index of the given CRE in the patient cohort. Pval represented at the top of each violin plot group is obtained by applying the Kruskal-Wallis chi-squared. The statistical test applied: Pairwise Wilcoxon rank-sum test. *= Pval< 0.05. Color legend of violin plot: Blue= Healthy samples; Green= Samples at Onset; Orange= Samples at Remission; Relapse= Samples at relapse. **d**) Left, qRT-PCR analysis of DCTD eRNA expression (eDCTD) or DCTD gene expression in B-ALL cells following CRISPR/Cas-9 with two different gRNAs (#1-#2) compared to a control gRNA. Relative fold changes were determined by the comparative treshold (ΔΔCt) method using β -actin as endogenous normalization control. Data are presented as mean ± SD of three independent experiments; middle, WB for DCTD in B-ALL cells to evaluate CRISPR/Cas-9 efficiency. β−actin was used as loading control; right, cell number analysis was performed in B-ALL cells treated as in D at different time points. Data are presented as mean ± SD of three independent experiments. **P≤ 0,01, ***P≤ 0,001 by Student’s *t*-test.

Altogether, these data provide evidence that DCTD expression and protein translation are driven by a productive enhancer placed 108Kb upstream of DCTD promoter, which is clonally amplified during cancer initiation and recurrence.

## DISCUSSION

While many principles of chromatin regulation have been elucidated in cultured cancer cells, epigenomic studies of primary cancer are uniquely valuable in capturing the genuine regulatory specificity of cancers. In this study, we profiled the cis-regulatory asset of a longitudinal cohort of clinically annotated patients to investigate the contribution of enhancers to the emergence and progression of BCP-ALL. We demonstrated that BCP-ALL phenotypical heterogeneity is sustained by engaging a plethora of previously unknown enhancers. We identified more than 120,000 active CREs, which contribute to cancer heterogeneity. More importantly, we observed dedifferentiation of cells independent of the molecular abnormalities, which ultimately converge to surge and repression of chromatin state specific to the BCP-ALL cancer stage. Indeed, we identified ∼11k stage-specific plastic CREs sustaining BCP-ALL phenotype. With a multi-omics integrative approach, we defined the role of these CREs and nominated long-range gene-regulatory interactions with the ultimate target genes, of which some, not surprisingly, are genes previously implicated in the BCP-ALL and hematopoietic malignancies such as EBF1, MYB, ETS1, and MYC and many others. Interestingly our data also uncovered specific driving transcription factors at the selected CREs and provided evidence of how the binding grammar converges to only a few key transcription factors, including EBF1, ETS1, ERG, and RUNX. On the other hand, we could identify enhancer-gene relationship at loci never implicated with lymphoid malignancy. That is the case with MYB and DCTD genes. Indeed, we experimentally demonstrated that the identified regions at the genes were dynamically modulated in dependency on the disease stage and, more importantly, are a key determinant of the target gene transcription and translational output.

The major obstacle to effective cancer therapy is the transcriptional and molecular heterogeneity, which stimulate the research of mechanisms able to act transversely across patients’ characteristics. Results obtained by our epigenetic study show that focus on the CRE may overcome the heterogeneity of the samples and the different causes of relapse in response to different therapies by individuating regulatory elements that are transversely active among the samples. The data generated in this study provide unique opportunities to characterize the landscape and functions of CREs across different BCP-ALL stages. We applied our selection by integrating eRNA data to identify the productive/activated enhancers. eRNAs are increasingly recognized to play important roles in the regulation of gene transcriptional circuitry in human cancers. We agree with the notion that eRNAs per se may serve as useful and highly precise therapeutic targets for future cancer intervention. This is particularly based on the high specificity of eRNA expression across tissues (64), and across cancer types (22,28) and provides a superior advantage to being a drug target as its inhibition will not affect other irrelevant tissues and, more importantly, does not completely abrogate the expression of target genes. In fact, with the aim to verify a regulatory interaction for the predicted peak-to-gene links, we applied CRISPR-Cas-9 editing to epigenetically inhibit the eRNA productive regulatory elements in BCP-ALL primary cell lines. Editing of the newly identified eRNA sequence controlling the Myb-HBSL1 complex resulted in strong inhibition of blast cell viability when compared with the effect exerted by control gRNA. In addition, we successfully achieved around 8-fold eDCTD downregulation by two different gRNAs, which led to strong downregulation of blast cell proliferation. Long-term observation of eDCTD-edited cells has provided evidence of a stable reduction in the proliferation rate that could facilitate the efficacy of chemotherapeutic agents.

In conclusion, this study supports the value of the epigenetic approach in identifying new tumorigenic elements that could be targeted only on the base of the cancer status. In particular, the identification of regulatory elements accessible and functionally relevant only in the relapsed phenotype could drive the development of new target therapy. Targeting cancer cell states rather than distinct genotypes may therefore represent a way to overcome the extensive genetic heterogeneity of cancer.

## MATERIALS AND METHODS

### Patients’ characteristics

Patients with newly diagnosed or relapsed BCP-ALL were treated at IRCCS Bambino Gesù Children’s Hospital (Rome, IT). The “newly diagnosed” group consisted of 26 patients (4 matched patients included), 19 males (73%), and 7 females (27%), with a median age at diagnosis of 7,5 years (range 0,4-18). The group of relapsed patients included 8 patients, 7 males (88%), and 1 female (12%), with a median age at diagnosis of 8,05 years (range 2-18) and a median age at relapse of 10.6 (range 4-23). Bone marrows (BMs) used as negative controls were obtained from age-matched or adult healthy donors (HBM) who donated BM for transplantation at Bambino Gesù Children’s Hospital. The Bambino Gesù Children’s Hospital Institutional Review Board approved the study (Prot n.495 del 11/04/2019).

### CD19-sorting selection

BMs derived from healthy donors and BCP-ALL patients were sorted by CD19 expression to select the B-cell compartment and blast cells, respectively. In brief, the whole BM samples were incubated with Human B RosetteSep (Stemcell Technologies, CAN) following the manufacturer ‘instructions.

### Cell lines, transfections and reagents

ALL-B cells were obtained by BCP-ALL bone marrow mononuclear cells infected with Epstein-Barr virus for immortalization (41). Both cell lines were cultured in RPMI-1640 medium (Euroclone) supplemented with 10% FBS (Thermo Fisher Scientific), 2mM glutamine (Thermo Fisher Scientific) and 40 µg/ml gentamicin. All cell lines were cultured at 37°C, in a humidified atmosphere with 5% CO_2_. Mycoplasma contamination was periodically checked by polymerase chain reaction (PCR) analysis, using the following primers:

Forward: 5’ –ACTCCTACGGGAGGCAGCAGTA- 3’

Reverse: 5’ –TCGACCATCTGTCACTCTGTTAAC- 3’

Nucleofection experiments of ALL-B cells were carried out using Amaxa 4D-Nucleofector X kit L (Lonza) according to the manufacturer’s instructions. Cells were analysed 36h after nucleofection by western blot (WB) or quantitative real-time PCR (qRT-PCR).

### CRISPR-Cas9 experiments

Lentiviral supernatants were generated into HEK293T cells by transient co-transfections of lentiviral CAS9-RFP Lenti Plasmid (Merck, USA) construct and appropriate amount of packaging vectors (Mission Lentiviral Packaging Mix, Sigma-Aldrich, USA) by TRANS-IT (TRANS-IT X2, Dynamic delivery System, Mirus, USA) following the manufacture’s protocol. After 48 hours supernatants were collected and employed to LAL-B cells in Retronectin (Takara Bionic Otsu, Shiga 520-2193, Japan) pre-coated no tissue culture 24-well plates (Falcon, BD, USA). Viral particles were centrifuged at 2000g, 90 minutes, at 32°C. Cells were harvested, washed with PBS (Pan Biotech, DH) and replated in RPMI (EuroClone, IT) with 10% FBS (Gibco, USA). After 48 hours, infected cells were sorted following RFP reporter signal to obtain a pure population. LAL-B/CAS-9 clone was transduced, following the Retronectin protocol, with lentiviral particles carrying single CRISPR guide RNA (Custom CRISPR gRNA Plasmid DNA, Merck, USA) directed to DCTD eRNA, -67 kb eRNA and -51 kb eRNA. CTRL-CRISPR gRNA was used as negative control.

### gRNAs sequences

**-67 kb cMyb**

5’-AAGAGGAAAAGGCGAGAAT-3’

5’-AGAGGAAAAGGCGAGAATC-3’

**-51 kb cMyb**

5’-TCATTGCTATATGTAGGTA-3’

5’-GCAAACGAAACACGACTCC-3’

**DCTD**

5’-ATACTCACGCCCGAGAGTC-3’

5’-AGGCTGCATCATCTTCAAA-3’

**CTRL**

5’- CGCGATAGCGCGAATATATATT-3’

**siRNA**

siRNA experiments of DCTD expression were performed by transfecting a specific pool of three double-stranded RNA oligonucleotides (siDCTD, cat. n. 1299003 – HSS102676, HSS1026777, HSS102678) or a control sequence (siControl, cat. n. 12935300) (Stealth, Thermo Fisher Scientific) using Amaxa 4D-Nucleofector X kit L (Lonza).

An antisense oligonucleotide targeting DCTD eRNA (CACGGAGCAUGGCAACCUGCAAACA) was purchased from Eurofins Genomics.

### Total cellular extracts and western blotting

Total cellular extracts were obtained as described in Bruno et al. (2006) (65). Proteins (25 µg) were separated by electrophoresis and transferred onto nitrocellulose membranes. After a blocking step in 5% nonfat-dried milk in 0.1% Tween-PBS, membranes were incubated with primary antibodies overnight at 4°C. After three washes in 0.1% Tween-PBS, membranes were incubated with the appropriate HRP-linked secondary antibodies (Bio-Rad) at room temperature for 45 min, washed with 0.1% Tween-PBS and analyzed by chemi-luminescence (GE Healthcare Life Science). Images were acquired and quantified using Alliance Mini HD6 system by UVITEC Ltd, Cambridge, equipped with UVI1D Software (UVITEC, 14-630275). The rabbit polyclonal antibodies used were: c-Myb (D2R4Y, Cell Signaling), DCTD (ab183607, Abcam), HBS1L (Proteintech). Mouse monoclonal antibody was b-actin (clone AC-15, Sigma-Aldrich).

### RNA isolation and quantitative real-time PCR

Total RNA was isolated from cells using EuroGOLD TriFast reagent (Euroclone) according to the manufacturer’s instructions. cDNA was synthesized from equal amount of RNA by reverse transcription using M-MLV reverse transcriptase (Thermo Fisher Scientific) and a mixture of random primers (Thermo Fisher Scientific). This single-stranded cDNA was then used to perform quantitative real-time PCR (qRT-PCR) with specific primers using PowerUP SYBR Green 2x Master Mix (Thermo Fisher Scientific) on a 7500 Fast Real-Time PCR System (Applied Biosystems), following the manufacturer’s instructions. Data were processed using the 7500 software v2.0.6 (Applied Biosystems). Relative fold changes were determined by the comparative threshold (ΔΔCt) method using β-actin as endogenous normalization control (66). Data are presented as mean ± SD of three independent experiments, performed in duplicate. Specific primers employed in qRT-PCR amplifications are listed in Table X. 11846609

### ChIP-sequencing

Chromatin immunoprecipitation (ChIP) assays were performed as previously described (Bruno T, Cancer Cell, 2002) by using anti-acetyl-Histone H3 (Lys27) antibody (Millipore). Equal amounts of precleared chromatin were added to antibody-bound Dynabeads (Thermo Fisher Scientific). About 6 ng of the immunoprecipitated chromatin was used to prepare the libraries for sequencing by following the manufacturer’s instructions (Swift Bioscences, Cat. No. 21024). The final libraries were controlled on an Agilent 2100 Bioanalyzer (Agilent Technologies) and sequenced in paired-end mode (2×75 bp) with NextSeq 500 (Illumina, CA).

### Promoter capture Hi-C

Hi-C in the B-ALL cell line was performed using the Arima-HiC Kit according to the manufacturer’s instructions. Briefly, 1 x 10^6^ cells were crosslinked with 1% formaldheyde, digested with a restriction enzyme cocktail, end-labeled with Biotin-14-dATP and then followed by ligation. The ligated chromatin was reverse cross-linked and sonicated using Bioruptor ultrasonicator to produce 300-500 bp fragments. Fragmented DNA was then size-selected to have a size distribution between 200-600 bp, and finally subjected to biotin enrichment. DNA libraries were prepared using Accel-NGS 2S Plus DNA Library Kit (Swift Bioscences, Cat. No. 21024), and the resulting libraries were amplified using the KAPA library amplification kit. Subsequently libraries were hybridized to specific SureSelect XT Human capture libraries (Agilent Technologies) and sequenced in paired-end mode (2×75 bp) with NextSeq 500 (Illumina, CA).

### RNA-sequencing

Total RNA was extracted from patient samples using Qiazol (Qiagen, IT), purified from DNA contamination through a DNase I (Qiagen, IT) digestion step and further enriched by Qiagen RNeasy columns for gene expression profiling (Qiagen, IT). Quantity and integrity of the extracted RNA were assessed by NanoDrop Spectrophotometer (NanoDrop Technologies, DE) and by Agilent 2100 Bioanalyzer (Agilent Technologies, CA), respectively. RNA libraries were generated using the same amount of RNA for each sample according to the Illumina TruSeq Stranded Total RNA kit with an initial ribosomal depletion step using Ribo Zero Gold (Illumina, CA). The libraries were quantified by qPCR and sequenced in paired-end mode (2×75 bp) with NextSeq 500 (Illumina, CA). For each sample generated by the Illumina platform, a pre-process step for quality control was performed to assess sequence data quality and to discard low-quality reads.

### ATAC- sequencing

To profile open chromatin, we used the ATAC-seq protocol developed by Buenrostro et al., (38), with minor modifications. Plasma cells were isolated from malignant and control bone marrow aspirates. 50,000 cells were washed once with 1X PBS and centrifuged at 500g for 5 minutes at 4°C. The cell pellet was lysed in ice-cold lysis buffer (10mM Tris-HCl pH 7.4, 10mM NaCl, 3mM MgCl_2_, 0.1% IGEPAL CA-630) to isolate the nuclei. If the cell pellet has been flash-frozen at −80°C, the morphology of the isolated nuclei was carefully inspected for integrity by trypan blue coloration. The nuclei were centrifuged at 500g for 5 minutes at 4°C and subsequently resuspended on ice in 50μl transposase reaction buffer containing 2.5μl of Tn5 transposase and 25μl of 2xTD buffer (Nextera DNA Sample preparation kit from Illumina). After incubation at 37°C for 30 minutes, the samples were purified with MiniElute PCR Purification Kit (Qiagen), eluting in 10µl elution buffer (10mM Tris-HCl pH 8). To amplify transposed DNA fragments, we used NEBNext High-Fidelity 2x PCR Master Mix (New England Labs) and the Customized Nextera PCR Primers. Libraries were purified by adding Agencourt Ampure XP (Beckman) magnetic beads (1:1 ratio) to remove remaining adapters (left side selection) and double purified (1:0.5 and 1:1.15 ratio) for right side selection. Libraries were controlled using a High Sensitivity DNA Kit on a Bioanalyzer (Agilent Technologies). Each library was then paired end sequenced (2×75bp) on a NextSeq 500 instrument (Illumina).

## COMPUTATIONAL METHODS

### ATAC-seq analysis and differential enrichment

The ATAC-sequencing reads quality was assessed with FastQC v0.11.9 (http://www.bioinformatics.babraham.ac.uk/projects/fastqc/). Reads were aligned to the reference genome hg19 using bowtie v2.3.5.1 (67) with default parameters. The conversion from sam to bam file was performed through view function of SAMtools v1.7 (68). BAM files were deduplicated with GATK v4.1.9.0 markDuplicates with default parameters. ATAC-seq peaks were called by MACS2 v2.2.6 with parameters *--format AUTO --nomodel --shift -100 --extsize 200 -B --SPMR --call-summit -q 0.01 -g hs*. BigWig (bw) files were obtained from the BedGraph (bdg) files with bedGraphToBigWig v4 (69) with default parameters. Finally, all peaks matching blacklisted regions (downloaded from https://www.encodeproject.org/files/ENCFF001TDO/@@download/ENCFF001TDO.bed.gz) were removed with the function *intersect* of bedtools suite v2.29.2 (70).

### Multidimensional scaling

Multidimensional scaling (MDS) was carried out on a normalized table in which each row is a peak of the master list and each column represent a different sample. The reads count was performed with bedtools *multicov* v2.29.2.

The reads count per peak was normalized through the R package *edgeR* v3.36.0 (71) and the obtained TMM was transformed through the log2(TMM+1) formula.

Finally, MDS was performed with the *cmdscale* function of *stats* R package v 4.1.2 applied at the samples distance matrix. The distance metric chosen is the following:

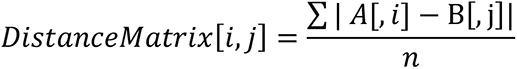

Both i and j are samples and n is the total number of peaks.

The scatterplot was performed on first-second and first-third components through ggplot2 R package v 3.3.5.

### Differential analysis of accessibility profiling of BCP-ALL

We first built a master list of the accessible regions identified in the profiling of each patient. Narrow peak files of each sample were concatenated, sorted and merged with bedtools *merge* function to finally obtain a list of all accessible region’s profiles in at least one patient. We then performed the differential analysis of samples by building a matrix of the peak read count of each sample on the master list. In order to accomplish the reads count the master list and all the samples’ bam were used as parameter of *bedtools multicov*. The count normalization was performed with the R package *edgeR* v3.36.0. Two comparisons were performed: Healthy vs Onset and Healthy vs Relapse. Peaks with differences in the means were extracted with *extractTest* edgeR function, and the p-value was corrected through the False Discovery Rate (FDR) method. Finally, the selected peaks whit FDR <= 0.0001 were divided into four groups (-log10(FDR) ≤ 1, −log10(FDR) ≤ 2, −log10(FDR) ≤ 3, −log10(FDR) ≤ 4) and visualized through a scatterplot performed with ggplot2 R package. To provide further significance to the differential analysis, we performed the randomization of the dataset by applying 100 random sample selections. At each iteration, we selected 17 samples randomly selected amongst our cohort of BCP-ALL samples (N=32). At each randomized set, we performed differential analysis as described above by divide the 17 selected sample into two groups: 6 Healthy and 11 Onset. Ontology analysis of the differential subset of peaks was performed by using GREAT v3.0.0 (72). Peaks annotation was performed on hg19 human genome and the basal plus extension was chosen with the following parameters: proximal 5.0 kb, upstream 5.0 kb and plus Distal up to 100.0 kb).

### Clonality and Penetrance index scoring strategy and relative analysis

The Clonality and Penetrance index are calculated by 2 independent analysis workflows which ultimately assigned to each genomic region included into the master list of accessibility the Clonality and Penetrance index. The RI is a patient specific standardized metric determined by the following formula calculated to the peak repertoire of each sample independently: Nscore= ((peak read count / peak size)⋅10^-6^))* 10^-3^ /total mapped reads (FPKM). The peaks’ read count are assessed with bedtools multicov function. Then, peaks in each sample are arranged from highest Nscore to lowest Nscore. The ordered list of Nscoring is divided into percentiles ranging from 1 (higher enrichment) to 100 (lower enrichment). All peaks present in the master list and absent in the given sample receive assigned RI=0. The penetrance index is assigned to each peak of the master list and represents the number of patients sharing each given peak in the patient cohort. At each peak is assigned an SI value from 1(only one patient carrying the given peak) to 32 (the totality of patients carry the given peak).

The relation between RI and SI was displayed through a boxplot obtained with the seaborn python library v0.11.2. Samples were divided into the relative disease group (Healthy, Onset, Remission, and Relapse), and the RI median value was calculated for each peak considering only peaks with RI different from 0.

The linear regression for each status (Healthy, Onset, Remission, and Relapse), the median Clonality index, and the Penetrance index data were fitted with the *sklearn LinearRegressor* model. Finally, the R^2^ was calculated on the fitted model. Moreover, a Linear Regressor and R^2^ were calculated, gathering all the data of each status. Both models and R^2^ were computed through an in-house python script relying on the Scikit-Learn python library v 1.0.2.

To assess the composition of cancer stages in each SI, we divided peaks from the master list into 32 groups (from SI=1 to SI=32). In each group, we calculate the percentage of peaks belonging to every disease group (Healthy, Onset, Remission, and Relapse). The stacked bar plot was used to display the results.

To assess the genomic distance from detected peaks and relative nearest TSS we divided the peaks into four lists, one for each disease group, containing only peaks detected in at least one sample within each group. We ran each list to the *annotatePeaks.pl* function included into HOMER suite (73). Then, peaks were divided into five groups of distance relative to the closest TSS (<5kb, 5kb-20kb, 20kb-100kb, >100kb). Finally, for each group was calculated the percentage contribution at each distance group and plot with an in-house R scripts. The HOMER results described above were also used to classify peaks according to the genomic context. Hence, peaks were divided into six classes (Non-Coding, Promoter, Exon, Intron, TTS, and UTR) according to the HOMER output.

### Observed and Expected (O/E) relationship of peaks at disease stage

We measured the significance of the observed peaks in our cohort and their relationship with the disease stage. A count of the observed number of peaks belonging to each Penetrance score was performed. The expected number of peaks for each status per SI was calculated applying this formula:

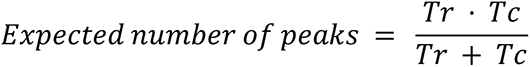

With:

Tr: total number of peaks in a row
Tc: total number of peaks in a column

Finally, we performed the observed-expected ratio for each disease group per SI.

### Assessment of peak dynamics during BCP-ALL evolution

Peaks were stratified according to their relative impact in driving cancer onset and relapse. We first assigned the median of the Clonality index and the Penetrance index to each peak of the master list. The final matrix was then populated with peak coordinates (hg19) at the rows and Clonality index median and penetrance index of each status at the columns (eight columns). The selection criteria to prioritize the phenotypical peak drivers are the following:

Peaks that harbor Onset Penetrance Index ≥ 9 and Healthy Penetrance Index ≤ 3 were selected. Peaks that show an increase of at least 20 in the difference between the Healthy Clonality index median and Onset Clonality index were selected. Finally, peaks that harbor an increase of at least 20 in the difference between the Remission Clonality index median and Relapse Healthy Clonality index were selected. Finally, we obtained a list of unique 11.077 unique peaks. Enrichment trends of the selected peak at each cancer stage were plotted as a boxplot with an in-house r script.

### Density distribution of phenotypically driver peaks

We measured the relationship of the selected peaks according to the cancer stage by calculating the density distribution of the penetrance score. We first calculated the ratio of each peak by the number of total samples at the disease stage and obtained a standardized score ranging between 0 to 1, where 0 is associated with a silent state and 1 to activation in all the patients in the disease stage cohort. Finally, we plotted the density distribution of peaks across the four-cancer stages through the *plot_density* function of the ggplot2 R package v 3.3.5.

### Heatmap of peak dynamics during BCP-ALL evolution

A heatmap was performed to assess the read count of the 11.077 selected peaks across all 32 samples. First, a read count of these sites was performed through the bedtools *multicov* v2.29.2 with standard parameters. The obtained reads count was normalized through the R package edgeR v. 3.36.0 in order to obtain the log2(TMM+1) metric. Finally, the TMMs were scaled, and the visualization was performed through an R script through the ComplexHeatmap R package v. 2.10.0. The cluster detection was accomplished according to the Ward2 hierarchical clustering to retrieve four final clusters classified as C1, C2, C3, and C4. The similarity across samples was evaluated through the Euclidean distance.

### Transcription Factors Identification

Peaks belonging to the selected four clusters above (C1, C2, C3, C4) were analyzed in order to infer the putative transcription factor able to bind towards these genomic regions. The HOMER findMotifsGenome.pl (parameter: --genome hg19) tool v4.11 was employed. For each of the significant transcription factors identified, we calculated the Observed/Expected ratio and displayed the data as a circular bar plot with an in-house r script.

### Integration with TCeA portal data

To infer the number of eRNA productive peaks, we integrated our peak selection with the TCeA portal (40 (https://bioinformatics.mdanderson.org/Supplements/Super_Enhancer/TCEA_website/) by downloading the Canonical Enhancer and Super-Enhancer data from available tumor types. We performed the intersection of the 2 datasets with our peak selection with bedtools intersect tools v2.29.2.

### Putative eRNA in the BCP-ALL cohort

In order to assess the amount of actively transcribed enhancers, we performed a bedtools intersect between the TCeA eRNA data and the blacklisted MACS2 output narrowpeak file of each patient. Finally, we plotted the MACS2 (74) *signalValue* of the actively transcribed peaks. The violin plot was ordered according to the median of the peaks’ enrichment in each patient.

### eRNA identification of RNA-seq BCP-ALL cohort

Total RNA sequencing was performed to estimate the eRNA expression across 8 samples (4 Healthy and 4 Onset) at the cancer phenotype’s selected regulatory regions. Quality control of the RNA-sequencing reads was obtained through FastQC v0.11.9. Reads were aligned to the reference genome hg19 using STAR (v. 2.7.9a) (75). The conversion from sam to bam file was performed through the view function of SAMtools v1.7.

C1 and C2 clusters (N=5216) were extended with the DHS sites downloaded from https://personal.broadinstitute.org/meuleman/reg2map/HoneyBadger_release/ to quantify the expression of the non-coding genomic regions across the 11 patient RNAseq. We then removed all regions intersecting exonic regions. The read count of the extended non-coding regions (N=4935) was evaluated through the multicov function of bedtools v2.29.2 with standard parameters.

The read count standardization was accomplished through the edgeR v. 3.36.0 in order to obtain the scaled log2(TMM+1) used for the heatmap visualization. The two distinct clusters were carried out by dividing the dendrogram generated from the hierarchical cluster into two clusters. Finally, the sites of the target genes were identified according to the nearest gene inferred by using the HOMER annotatePeaks.pl (parameter: --genome hg19) function v4.11.

### Promoter-Capture Analysis

Paired end FASTQ files were aligned against the hg19 genome. The KR-normalized contact matrices and the loop annotation were performed with Juicer (76) v 1.9.9. Loops were called at 5kb, 10kb, and 25kb resolution.

We classified each promoter-CRE looping by intersecting the bedpe file generated by Juicer and the promoter coordinates (downloaded from https://egg2.wustl.edu/roadmap/data/byDataType/dnase/BED_files_prom/regions_prom_E00 1.bed) using bedtools intersect v2.29.2. We calculated the number of interactions in three different target-anchor loops (promoter-promoter, promoter-NonCoding, and NonCoding-NonCoding). Moreover, we assessed the looping length in genomic coordinates according to the distance from anchor to targets into five classes (<50kb, 50kb-200kb, 200kb-500kb, 500kb-1Mb, and >1Mb).

### RNA-seq analysis of LAL-B and B cells during differentiation

Differential expression analysis across LAL-B cell line and normal B cells (77) (GEO : GSE118165 ; Table 1). Samples were aligned to hg19 through STAR and quantified with RSEM (78). Both the alignment and quantification were performed through the nf-core/rnaseq v3.0 from NEXTFLOW v21.05.0-edg (79) with parameters: --aligner star_rsem. The quantification files were normalized through the standard edgeR (v 3.36.0) pipeline. Once the normalization step was carried out, we selected only the target gene of the selected enhancers and compute both the Ward2 hierarchical clustering and the heatmap with the R package ComplexHeatmap v 2.10.0.

Identification of the enhancer-target genes was manually curated by integrating ENCODE data (80) (ChIAP-pet, H3k27ac ChIP-seq), RNA-seq from TCGA and Promoter-Capture of LAL-B.

### ATAC-seq of B cells at multiple differentiation states

B-cell ATAC-seq FASTQ files were available at the GEO GSE118189 (Table 3).

The FASTQ quality was assessed with FastQC v0.11.9. Reads were aligned to the reference genome hg19 using bowtie v2.3.5.1 with default parameters. The conversion from sam to bam file was performed through view function of SAMtools v1.7. BAM files were deduplicated with GATK v4.1.9.0 markDuplicates with default parameters. ATAC-seq peaks were called by MACS2 v2.2.6 with parameters --format AUTO --nomodel --shift -100 --extsize 200 -B -- SPMR --call-summit -q 0.01 -g hs. BigWig (bw) files were obtained from the BedGraph (bdg) files with bedGraphToBigWig v4 with default parameters. Finally, all peaks matching blacklisted regions (downloaded from https://www.encodeproject.org/files/ENCFF001TDO/@@download/ENCFF001TDO.bed.gz) were removed with the function intersect of bedtools v2.29.2.

An assessment of read count both for LAL-B cell line, and the normal B-cell was performed through the bedtools multicov function v2.29.2. Finally, the read count file was standardized according to the library size through the edgeR R package v 3.36.0 and plotted through the ComplexHeatmap R package v 2.10.0. Both rows and columns of the resulting heatmap were clustered through the Ward2 unsupervised hierarchical clustering methods computing the distance through the Euclidean distance.

### Assessment of selected CRE clonality during BCP-ALL evolution

The clonality trends of the selected enhancer were assessed by analyzing Clonality Index across the four cancer stages and depicted as violin plots.

For each enhancer, the Clonality indexes were plotted, and statistical tests were performed: Kruskal-Wallis rank-sum test followed by Dunn’s Test.

### Analysis of Transcription Factors (TF)

We inferred the Transcription factor binding to the selected 111 enhancer, by downloading 32 ChIPseq of Transcription Factors: AP1: 8 samples, ATF3 : 4 samples, EBF1 : 2 samples, ELK4 : 2 samples, ERG:1 sample, ETS1 : 2 samples, ETV4 : 3 samples, FRA2 : 2 samples, RUNX1 : 5 samples, RUNX2 : 2 samples. Data were available in fastq format from the ChIPAtlas (81) (all the downloaded files annotation are gathered in the table 5). The fastq alignment and ChIP-seq seq peak detection were performed as described in the previous sections. For each transcription factor was performed the intersection between the selected enhancer and the blacklisted files with bedtools intersect v2.29.2. Finally, we ranked the TF by the number of the relative signal matching the selected enhancer.

### Pan-cancer analysis of H3K27ac ChIP-seq at selected 130 CREs

We downloaded all the available cell lines and primary cancer cell lines profiled for H3K27ac ChIP-seq available in ENCODE (all the downloaded files annotation are gathered in the table 6). Peak profiles of each BAM file were obtained as the follows: i) sorted with samtools sort v1.7; ii) the duplicated reads were removed with GATK v4.1.9.0 markDuplicates with default parameters; iii) the resulting file was indexed through samtools index v1.7; v) peaks were called through the MACS2 v2.2. callpeak function (parameters: --format AUTO -B --SPMR --call-summits -q 0.01); vi) BigWig (bw) files were obtained from the BedGraph (bdg) files with bedGraphToBigWig v 4 with default parameters. We assigned the Clonality Index to each significant peak as previously described. Clonality index annotation at the selected 130 CREs were shown as a heatmap where rows were arranged to display the enhancer detected in the highest number of cell lines at the top. Moreover, Ward2 hierarchical clustering was performed on the column and the sample distance was computed with the Euclidean distance.

### Gene dependency analysis

The fitness scores (CHRONOS) describing the effect caused by CRISPR knockout of 17.393 genes were downloaded from DepMap Public 21Q3 portal, table name CRISPR_gene_effect.csv. The data were restricted to only genes that resulted up-regulated from the differential analysis between LALB and B cells (N=106) and then plotted CHRONOS score of all available cell lines. The procedure to determine this list of genes was previously described. We further selected the genes exhibiting strong dependency specific to only BCP-ALL cell lines by ranking each CHRONOS score x disease cell lines. Only the genes showing lower CHRONOS score (high dependency) to BCP-LL cell lines (697, JM1, SEM, RCHACV, NALM6, REH, ROS50, SEMK2, HB1119, NALM16, P30OHK) were retained and plotted with an in-house R script.

### Data availability

The raw sequencing data are available on the ENA portal at accession number PRJEB52642.

## Acknowledgments

We want to acknowledge and thank all patients and their families for the support and for donating the research samples. This work was supported by the Italian Association for Cancer Research (A.I.R.C.) 15255 and MFAG-20096.

G. Corleone has received funding from AIRC and from the European Union’s Horizon 2020 research and innovation program under the Marie Skłodowska-Curie grant agreement No. 800924.

## Author contributions

All the authors discussed the experiments and contributed to the text of the manuscript. C.S. and M.C. performed most of the experiments and analyzed data. F.D.N. performed NGS experiments. G.C. performed the bioinformatics and statistical analyses and wrote the manuscript. S.D.G. and C.C. performed the bioinformatics and statistical analyses. V.B. and A.P. provided patient material. F.L. supervised and supported the study, wrote the manuscript, V.F. and M.F. conceived the project, obtained funding, supervised the experiments, and wrote the manuscript.

## Conflict of interest disclosure

The authors declare no competing financial interest.

## SUPPLEMENTARY FIGURES

**Supplementary Figure 1.**

**a)** Stacked bar chart depicting the percentage of ATAC-seq peaks x disease stage (Healthy- Onset- Remission-Relapse) in the function of the genomic distance to the closest TSS. Color legend: Blue= Healthy samples; Green= Samples at Onset; Orange= Samples at Remission; Relapse= Samples at relapse. **b)** Stacked bar chart showing the absolute number of ATAC-seq peaks at each given genomic annotation in Healthy, Onset, Remission, and Relapse groups. Color gradient from Purple to Green: Non-coding, Promoter, Exon, Intron, TTS, UTR. Annotation generated with HOMER suite. **c)** Upset plot of detected peaks at TSS (Promoter-like) proximity among the different groups of patients. X-axis: Intersection combination; Y-axis: the absolute number of detected sites. Color legend: Blue= Healthy samples; Green= Samples at Onset; Orange= Samples at Remission; Relapse= Samples at Relapse; Violet: Barchart of the number of detected sites at each intersection. **d)** MA plot of the differential peak accessibility. Left: Remission vs. Onset; Middle: Remission vs. Relapse; Right: Relapse vs. onset. X-axis: Log2(Peak mean), Y-axis: Log Fold Change of the differential accessibility of peaks. N= number of significant differential peaks identified in the analysis where logFC>0.7 is upregulation in Healthy; logFC<-0.7 is upregulation respectively at Onset (green), Remission (orange), and Relapse (red). **e**) Upset plot of differential peaks among all the differential analyses performed. X-axis: Intersection combination; Y-axis: the absolute number of detected sites. Color legend: Blue= Healthy samples; Green= Samples at Onset; Orange= Samples at Remission; Relapse= Samples at relapse. **f**) Ontologies of the differential accessible sites at Healthy vs. Onset (left) and Healthy vs. Relapse right. Top: Biological Process; Middle: Disease Ontology; Bottom: Mouse Phenotype. The analysis is performed with the GREAT tool. Color scheme: Green= Upregulation at Onset; Red= upregulation at Remission **g)** Upset plot of differential peaks among all the differential analyses performed. X-axis: Intersection combination; Y-axis: the absolute number of detected sites. Color legend: Blue= Healthy samples; Green= Samples at Onset; Orange= Samples at Remission; Relapse= Samples at relapse. **H)** Peak signal normalized across the full cohort of patients at selected genomic windows. Color legend: Blue= Healthy samples; Green= Samples at Onset; Orange= Samples at Remission; Relapse= Samples at relapse. Clonal CREs at the Onset and Relapse were further selected.

**Supplementary Figure 2**

**a)** Line plot of the linear regression between clonality index and the penetrance index among the four disease groups. y-axis = Clonality index, x-axis = Penetrance index. The coefficient of determination of each given linear analysis performed is shown on the bottom right. Color legend: Blue= Healthy samples; Green= Samples at Onset; Orange= Samples at Remission; Red= Samples at relapse.

**Supplementary Figure 3**

**a)** Density distribution of the selected 11k (C1, C2, C3, C4) CREs in healthy, onset (top) and remission and relapse (bottom). Y-axis= CREs density; x-axis= intra group normalized penetrance score. Color legend: Blue= Healthy samples; Green= Samples at Onset ; Orange= Samples at Remission; Red= Samples at relapse. **b)** Barchart shows the number of the selected peak in relationship with the genomic distance to the closest TSS. **c)** Gene ontology analysis of the selected peaks at three different databases: Biological process (blue); Disease ontology (red); Human Phenotype (violet). Significance is depicted on the x-axis (FDR) **d)** Violin plot shows peak enrichment over background at eRNA loci in each sample identified by integrating the TCeA portal. Samples are sorted from low (left) to high (right) by the median of peak enrichment. **e)** Upset plot shows the common peaks between the master list of onset, relapse, and LAL-B cell lines. Color legend: Green= Samples at Onset ; Red= Samples at relapse; Violet= LAL-B. **f)** Dot plot shows the frequency (y-axis) of the Transcription Factors at the 108 selected CRE. Transcription factors are ranked by the number of binding events among the selected 108 CREs.

**Supplementary Figure 4**

**a)** Heatmap shows the clonality score of the selected CREs (N=118) from H3k27ac ChIP-seq available on 159 cell lines (ENCODE data). Data were gathered by unsupervised clustering at the column and supervised at the rows. Column: Cell Lines Rows: CREs are named by the closest gene. Color legend: Blue= lower clonality index, Red= higher clonality index; White= no signal.

**Supplementary Figure 5**

**a)** Example of Promoter-Capture HiC looping in LAL-B (top) together with signals of (from top to bottom) ATAC-seq of LAL-B, CTCF ChiaPet of K562, Pol2 ChiaPet of K562, ATAC-seq from patients at Healthy, Onset, Remission and Relapse. The genomic window covers the coordinates of MYC locus (chr8:127,025,953-130,946,417), reference genome is HG19. Color legend: Blue= Healthy samples; Green= Samples at Onset; Orange= Samples at Remission; Relapse= Samples at relapse **b)** The heatmap depicts the unsupervised clutering of ATAC-seq data at the C1, C2, C3, C4 selected loci in Naïve-B cell (light grey), Memory B-cell (sky-blue), Bulk Bcell (dark-grey) and LAL-B (violet). Enrichment data are log2(Z-scaled). Color legend: Blue= low enrichment, Red= high enrichment. **c)** Dot plot showing the Chronos score (x-axis) of the selected genes (N=106 genes) in all the vailable cell lines (grey dots). ALL-B cell lines: 697, JM1, SEM, RCHACV, NALM6, REH, ROS50, SEMK2, HB1119, NALM16, P30OHK are colored violet.

**Supplementary Figure 6**

**a)** Bar Chart shows the cumulative enrichment of gene expression of single-cell types of the most expressing MYB gene. Colors depicts cell groups as follows: Red= Blood and immune cell types; Light blue= Undifferentiated cells; Blue= Glandular epithelial cells; Purple= endothelial cells; Orange= Pigment cells. Data obtained from Human Protein Atlas portal https://www.proteinatlas.org/. **b)** Box plot shows RNA expression of MYB in cancer tissues from TCGA. Data obtained from Human Protein Atlas portal https://www.proteinatlas.org/. **c)** MYB mRNA levels were analyzed by qRT-PCR in bone marrows samples of healthy donors (4), or from B-ALL patients at the time of onset (3), remission (3) or relapse (3). Relative fold changes were determined by the comparative threshold method (ΔΔCt) using β-actin as endogenous normalization control. Data are presented as mean ± SD of three independent experiments. ***P ≤0.001. **d)** Cumulative single-cell ATAC-seq signal in healthy Plasma cells (TOP) and Memory B Cells (Bottom) at MYB/HBS1L genomic window. Data obtained from http://catlas.org/humanenhancer/#!/. **e)** Boxplots show the Ratio between peaks (N-score) at enhancer at 51kb (left) and enhancer at 67kb (right) and MYB promoter signal in the patient cohort. Each dot represents a patient. Color legend: Blue= Healthy samples; Green= Samples at Onset; Orange= Samples at Remission; Relapse= Samples at relapse. Statistical tests performed: Kruskal-Wallis rank-sum test followed by Dunn’s Test. *P≤ 0.05. **f)** qRT-PCR analysis for Myb (left) or HBS1L (right) expression performed in B-ALL cells following CRISPR/Cas-9 of −51 kb region or −67 kb region using in each two different gRNAs (#1-#2), compared to a control gRNA. Values were normalized with β-actin mRNA levels using ΔΔCt method. Data are presented as mean ± SD of three independent experiments. ***P ≤0.001, **P ≤0.01 by Student’s *t*-test.

**Supplementary Figure 7**

**a)** Bar Chart shows the cumulative enrichment of single cell gene expression of different cell types of the most expressing DCTD gene. Colors depicts cell groups are showed on the right legend. Data obtained from Human Protein Atlas portal https://www.proteinatlas.org/. **b)** Protein expression of DCTD. Percentage of patients (y-axis) with high and medium DCTD protein level in different cancer cell types. Color code of the barchart is according to the type of normal organ the cancer originates. Data obtained from Human Protein Atlas portal https://www.proteinatlas.org/. **c)** Gene expression of DCTD gene (y-axis)in cancer (red) vs normal (green) tissues in cancer types available on TCGA portal. Cancer types exhibiting significance difference are colored red on the cancer type labeL (top). Data obtained from GEPIA2 (http://gepia2.cancer-pku.cn/). **d)** DCTD mRNA levels were analyzed by qRT-PCR in bone marrows samples of healthy donors (4), or from B-ALL patients at the time of onset (3), remission (3) or relapse (3). Values were normalized with b-actin mRNA levels using 1′1′Ct method. Data are presented as mean ± SD of three independent experiments. ***P ≤0.001, **P ≤0.01 **e)** Left, WB analysis of total cellular extracts from B-ALL cells transfected with siRNA oligonucleotides targeting DCTD (siDCTD) or a control sequence (siControl). b-actin was used as loading control. The same B-ALL cells were analyzed for cell number (middle) and for DCTD mRNA levels by qRT-PCR (right). Data are presented as mean ± SD of three independent experiments. ***P ≤0.001, **P ≤0.01. **f)** RNA expression data as normalized transcript per million of reads (y-axis) in cell lines. Color codes are according to the tissue of origin of the cell lines. Data obtained from Human Protein Atlas portal https://www.proteinatlas.org/. **g)** Boxplots show the Ratio between peaks (N-score) at the selected enhancer and the MYB promoter signal in the patient cohort. Each dot represents a patient. Color legend: Blue= Healthy samples; Green= Samples at Onset; Orange= Samples at Remission; Relapse= Samples at relapse. Statistical tests performed: Kruskal-Wallis rank-sum test followed by Dunn’s Test. *P≤ 0.05.

---

|  | Newly Diagnosed Patients | Relapsed Patients |
| --- | --- | --- |
|  | N.or median (% or range) | N.or median (% or range) |
| Number Of Patients | 26 (4 matched) | 8 |
| Gender |  |  |
| Females | 7 (27%) | 1 (12%) |
| Males | 19 (73%) | 7(88%) |
| Age at diagnosis (years) | 7,5 (0,4 - 18) | 8,5 (2 - 18) |
| Age at Relapse (years) | - | 10,6 (4 - 23) |
| Disease Status |  |  |
| Diagnosis | 26 (100%) | - |
| 1^ Relapse | - | 2 (25%) |
| 2^ Relapse | - | 5 (62%) |
| 3^ Relapse | - | 1 (12%) |
| Molecular abnormalities |  |  |
| None | 14 (64%) | 2 (25%) |
| t(12;21) (TEL/AML1) | 4 (18%) | 1 (12%) |
| t(9;22) (BCR/ABL) | 1 (4%) | 1 (12%) |
| t(9;11) | 1 (4%) | 0 (0%) |
| t(3;9) | 1 (4%) | 0 (0%) |
| t(3;20) | 1 (4%) | 0 (0%) |
| t(9;10) | 1 (4%) | 0 (0%) |
| t(5;11) | 0 (0%) | 1 (12%) |
| t(6;9) | 0 (0%) | 1 (12%) |
| t(7;9) | 0 (0%) | 1 (12%) |
| t(11;13) | 0 (0%) | 1 (12%) |
| t(16;1) | 0 (0%) | 1 (12%) |
| t(8;14) | 0 (0%) | 1 (12%) |
| r(KMT2A) | 1 (4%) | 1 (12%) |
| Aneuploidy |  |  |
| Yes | 8 (36%) | 5 (62%) |
| No | 14 (64%) | 3 (37%) |
| *Defined as chromosome number > or < 46 |  |  |

## Notes

### Competing Interest Statement

The authors have declared no competing interest.

