## Supplementary figures and images for "Enhancer plasticity sustains oncogenic transformation and progression of B-Cell Acute Lymphoblastic leukemia"

### Extended data 1

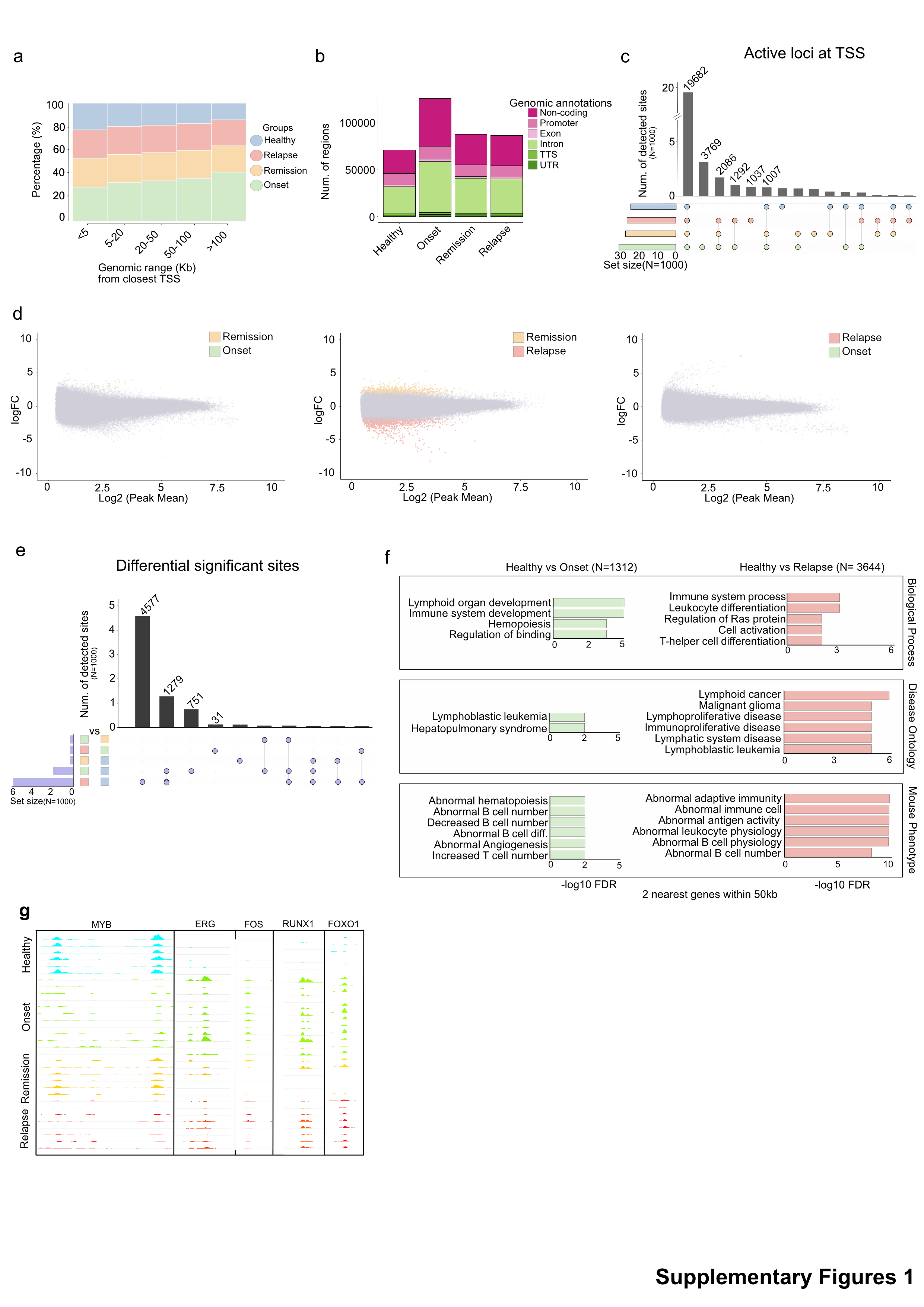

### Extended data 2

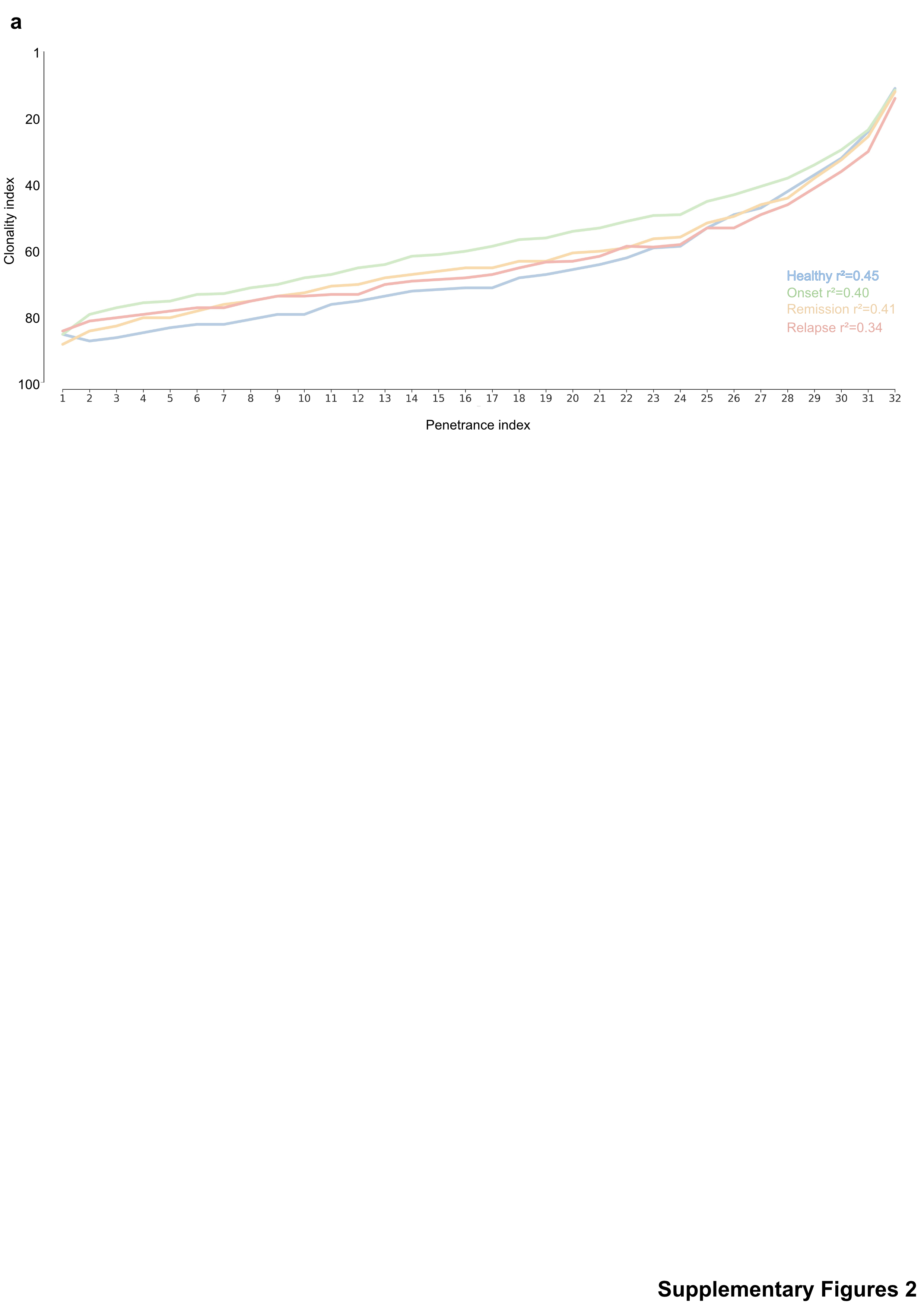

### Extended data 3

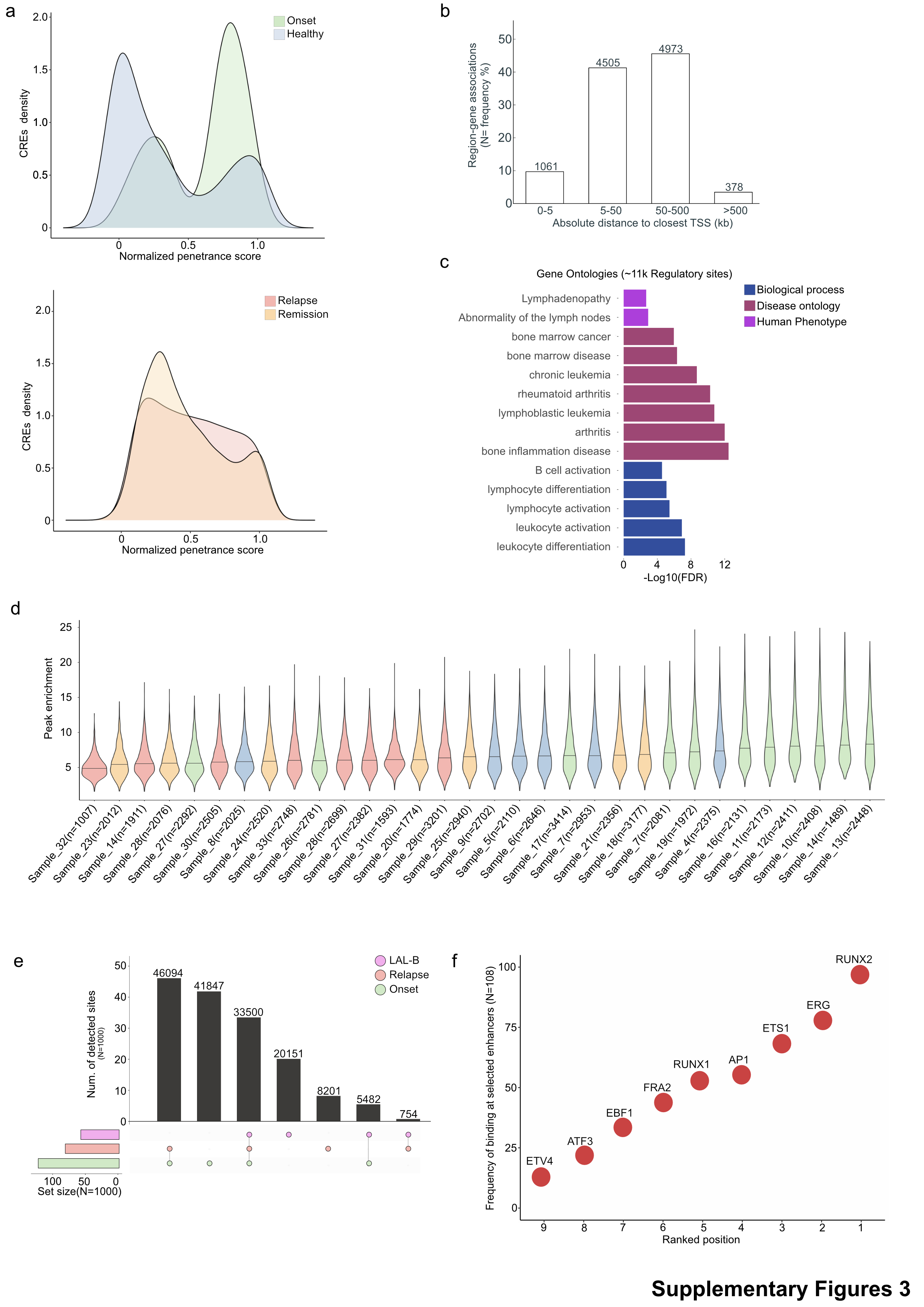

### Extended data 4

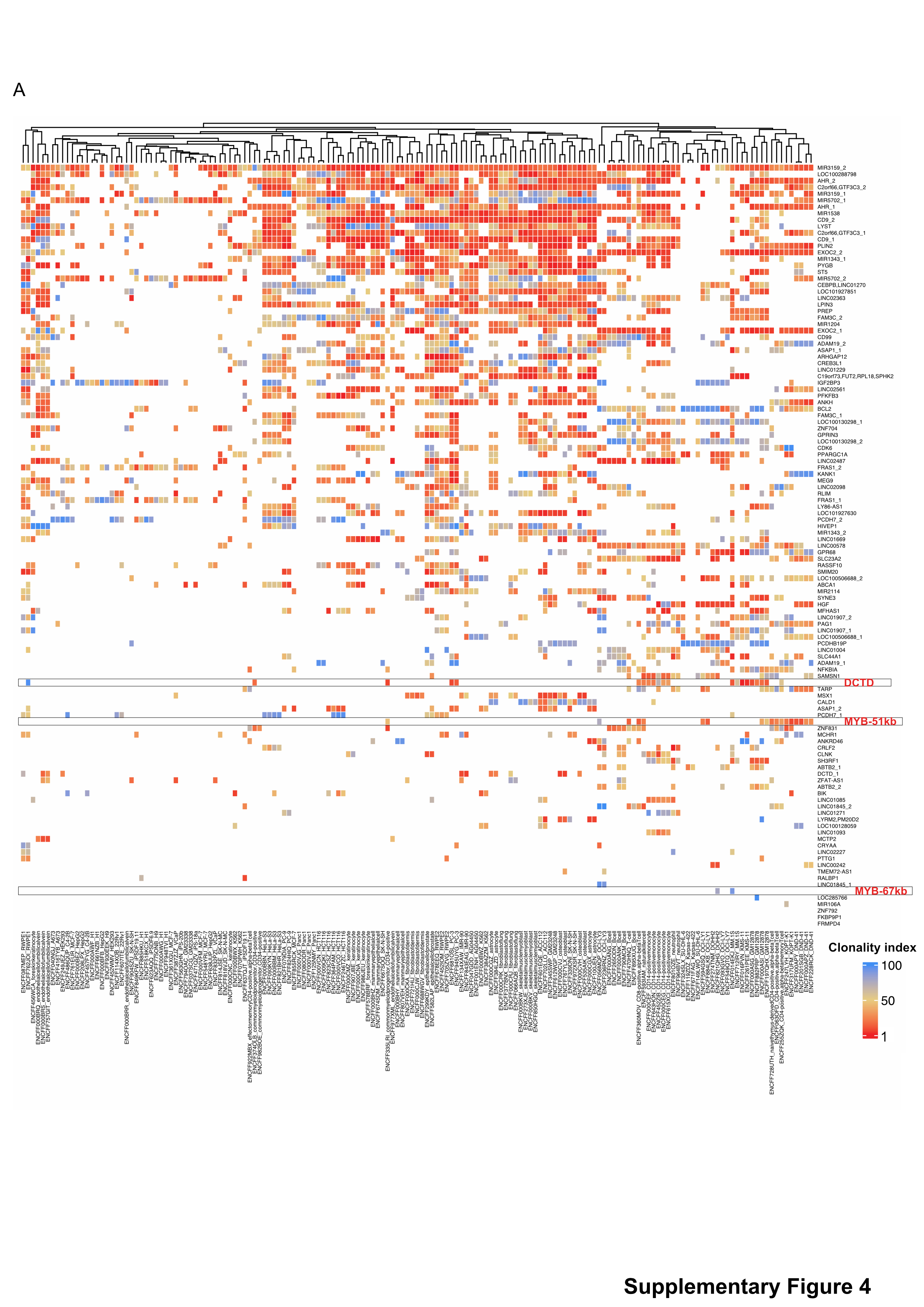

### Extended data 5

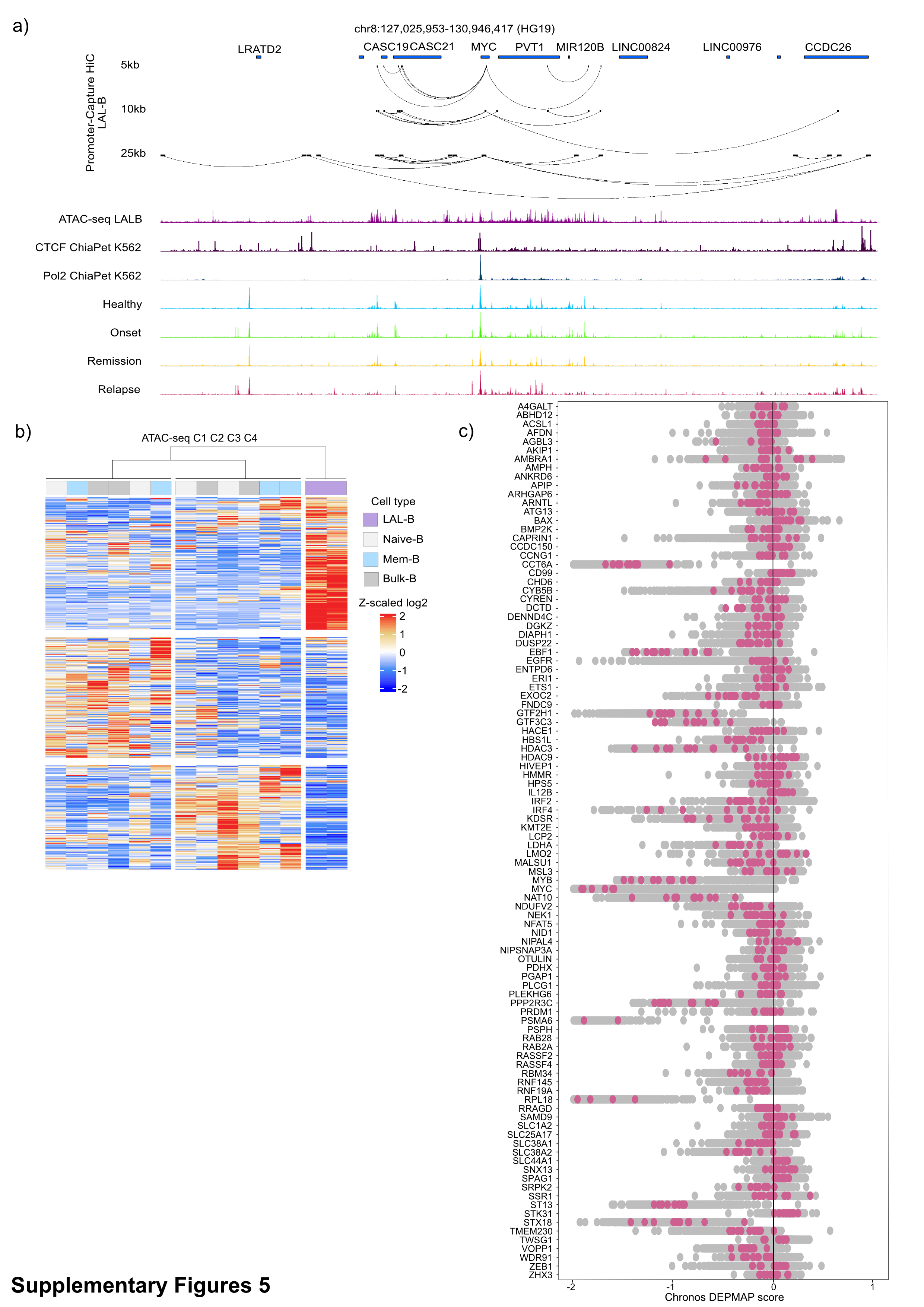

### Extended data 6

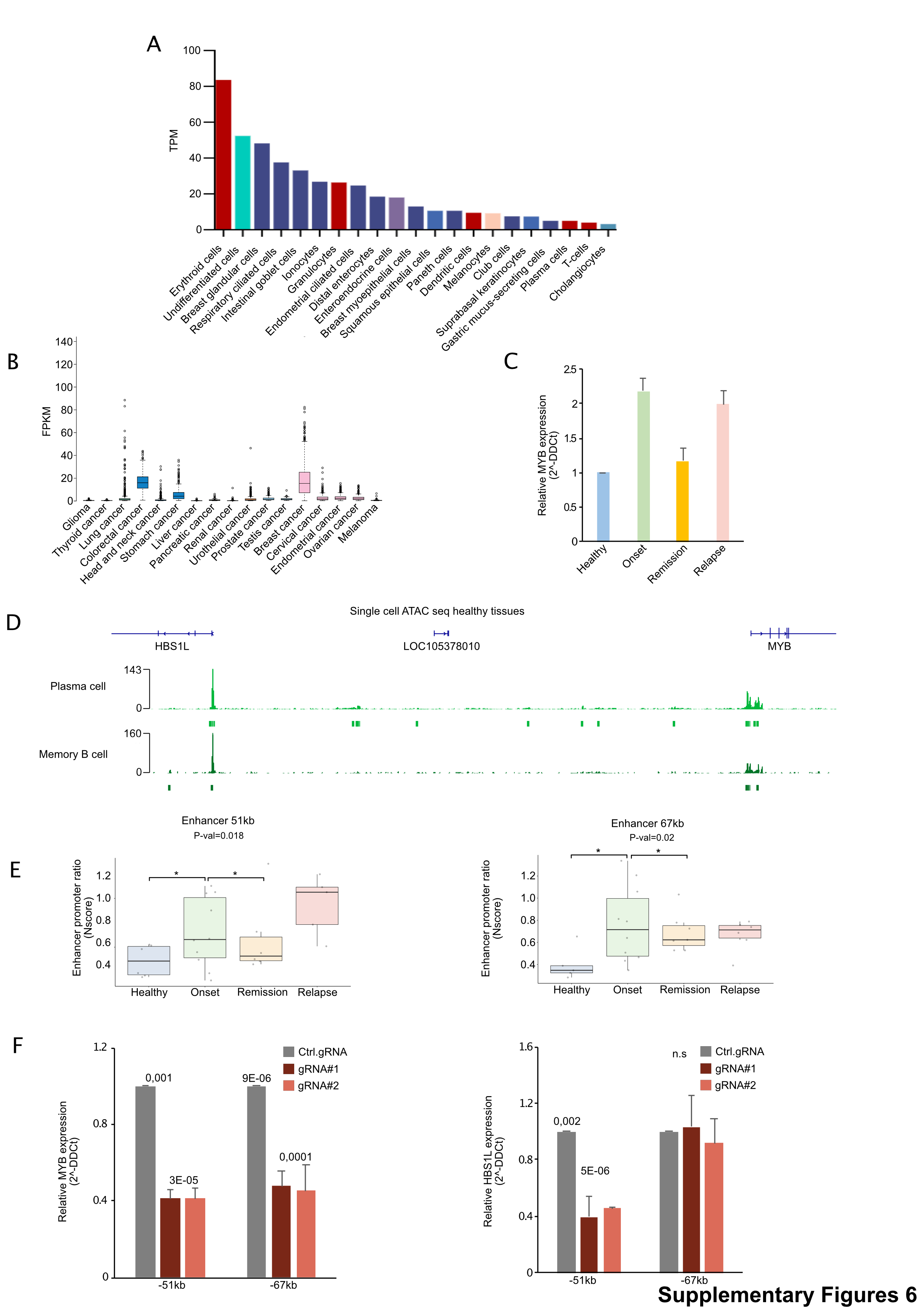

### Extended data 7

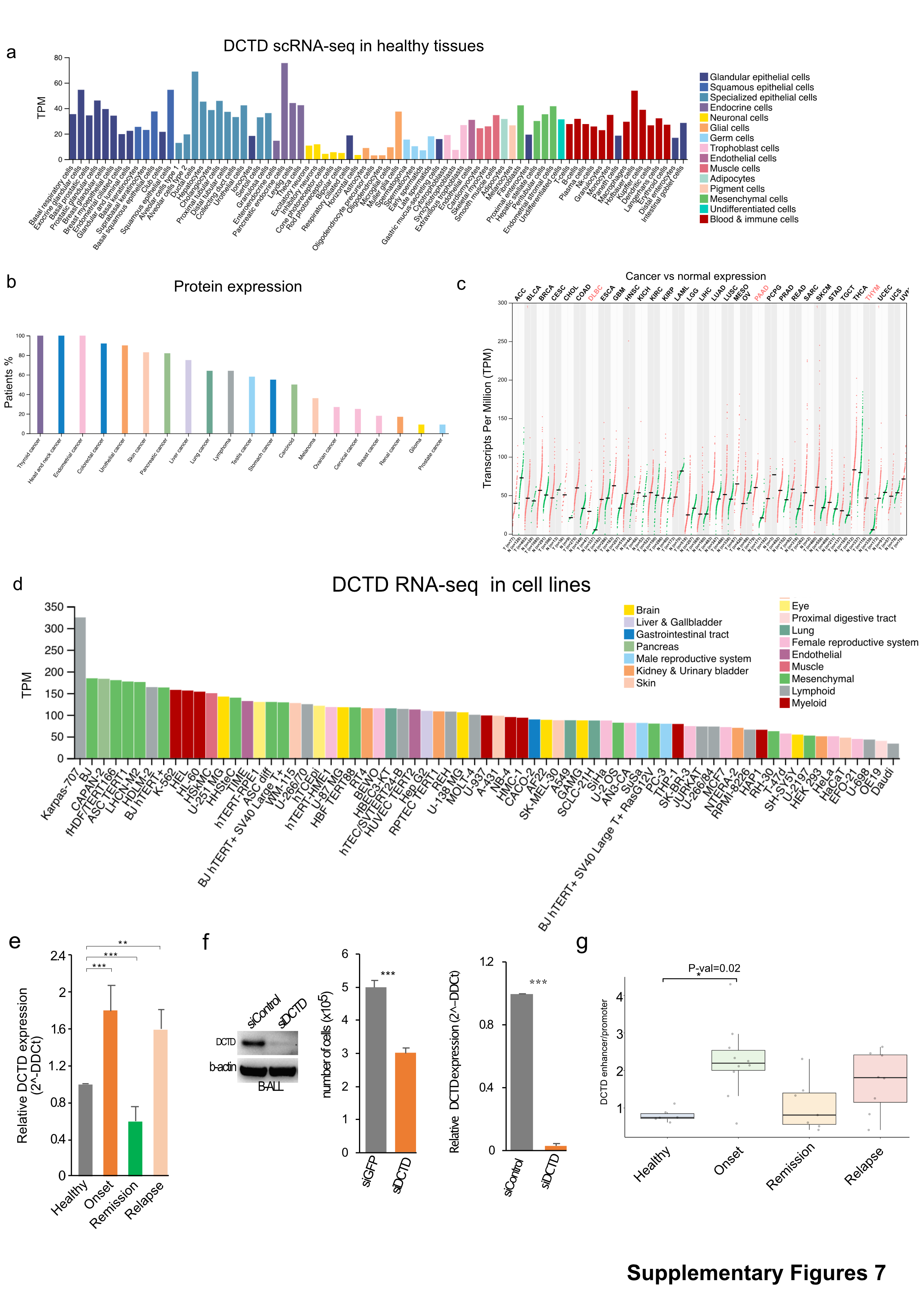
